# Acentrosomal spindles assemble from branching microtubule nucleation near chromosomes

**DOI:** 10.1101/2022.02.28.482415

**Authors:** Bernardo Gouveia, Sagar U. Setru, Matthew R. King, Howard A. Stone, Joshua W. Shaevitz, Sabine Petry

## Abstract

Microtubules are generated at centrosomes, chromosomes, and within spindles during cell division. Whereas microtubule nucleation at the centrosome is well characterized, much remains unknown about where, when, and how microtubules are nucleated at chromosomes. To address these questions, we reconstituted microtubule nucleation from purified chromosomes in meiotic *Xenopus* egg extract and found that chromosomes alone can form spindles. We visualized microtubule nucleation at chromosomes using total internal reflection fluorescence microscopy to find that this occurs through branching microtubule nucleation. The initial branches nucleate near and towards kinetochores, helping explain how kinetochores might be efficiently captured. By depleting molecular motors, we find that the organization of the resultant polar branched networks is consistent with a theoretical model where the effectors for branching nucleation are released by chromosomes, forming a concentration gradient around them that spatially biases branching nucleation. In the presence of motors, these branched networks are organized into multipolar spindles.

## INTRODUCTION

Microtubules originate from centrosomes, chromosomes, and spindle microtubules in dividing eukaryotic cells to form mitotic and meiotic spindles. Chromosomal microtubule generation is particularly critical in cells that do not contain centrosomes, such as plant cells and meiotic egg cells in animals^1–4^. While microtubule nucleation from centrosomes has been well studied, it remains poorly understood how microtubules are generated around chromosomes because spindle microtubules cannot be resolved from one another, nor can their exact origins be determined in cells. Thus, it remains a fundamental question in cell biology to understand where, when, and how microtubules are nucleated at chromosomes to build a spindle that successfully captures and segregates chromosomes during cell division.

Experiments in meiotic *Xenopus* egg extracts demonstrated that chromatin alone can generate spindles and tune a spindle’s size and shape^5–7^. Though revelatory, in lieu of actual chromosomes these studies used bacterial DNA on beads to form chromatin beads, which have different shapes and different amounts of chromatin compared to chromosomes. They also lack kinetochores, the landing pads on chromosomes where microtubules that make up kinetochore fibers bind before pulling sister chromatids apart. Indeed, experiments in cultured mitotic cells have shown that microtubules also form in the vicinity of kinetochores during prometaphase^8–15^, and in later stages at kinetochores on chromosomes unattached to existing spindle microtubules as well as far away from the spindle equator, suggesting a back-up mechanism to capture unaligned chromosomes^11,12^. Microtubule nucleation near kinetochores has not been observed directly in a meiotic system, where this is perhaps even more important than in mitosis given the lack of centrosomes that have traditionally been considered the origin of kinetochore fibers.

One way to generate microtubules around chromosomes is via the RanGTP gradient^16^. The chromatin-bound guanine nucleotide exchange factor RCC1 equips Ran with GTP. RanGTP then releases spindle assembly factors (SAFs) sequestered by importin proteins, allowing them to promote microtubule nucleation around chromosomes, as has been shown in *Xenopus* egg extracts using chromatin beads^17^ or sperm chromatin^18–22^, and in mitotic cells^23^. Cytoplasmic RanGAP1 deactivates RanGTP in a spatially uniform manner. This combination of a localized source and homogeneous degradation creates RanGTP and SAF gradients centered at chromosomes.

A critical SAF is the protein TPX2^24^, which, along with the protein complex augmin, the *γ*-tubulin ring complex (γ-TuRC), and XMAP215, promotes nucleation of new microtubules from the surface of existing microtubules^25^. This process, known as branching microtubule nucleation, generates the majority of spindle microtubules in mitotic cells^26^ and *Xenopus* egg extracts^27^. Yet, where exactly branching microtubule nucleation occurs around chromosomes remains an open question.

There could be other microtubule nucleation pathways at play that originate from chromosomes. For example, the chromosomal passenger complex (CPC) at the centromeres of chromosomes^28^ promotes microtubule growth by inhibiting the microtubule depolymerase MCAK, thereby promoting microtubule polymerization in *Xenopus* egg extracts^29^. It was shown that spindles can assemble around sperm nuclei in a manner independent of RanGTP but that requires the CPC^30^. The degree to which chromosomal microtubule nucleation differentially depends on RanGTP and the CPC, and whether branching microtubule nucleation occurs, remain to be determined.

Most quantitative studies to date have focused only on how microtubule nucleation sustains the steady-state metaphase spindle^27,31–33^, but do not address the question of how a spindle builds its microtubule network starting from scratch at the end of interphase. Similarly, simulations have focused on either the steady-state metaphase spindle^31^ or how centrosomes search for and capture kinetochores^34,35^. Currently, no model exists to describe how microtubules nucleate from chromosomes in the early stages of spindle assembly.

To explore these questions, we combine experiments that reconstitute microtubule nucleation from purified chromosomes in meiotic *Xenopus* egg extract with a mathematical model of branching nucleation in a SAF gradient. By directly visualizing microtubule nucleation at chromosomes using total internal reflection fluorescence microscopy (TIRFM), we reveal where individual microtubules nucleate, at what distances and orientations with respect to kinetochores they form, and how this is influenced by the amount of chromatin and number of chromosomes. By comparing experimental results to our model, we find that RanGTP-mediated branching microtubule nucleation in the vicinity of chromosomes provides the main source of microtubules in acentrosomal spindles.

## RESULTS

### Branching microtubule nucleation is initiated near chromosomes

It is challenging to directly visualize the nucleation of individual microtubules near chromosomes in cells because the spindle becomes dense with microtubules within minutes^7,36,37^. To overcome this obstacle, we reconstituted chromosomal microtubule nucleation *ex vivo*. Briefly, we purified mitotic chromosomes with fluorescent kinetochores from cultured mitotic HeLa cells. To enable live visualization of individual microtubules nucleating and growing at the onset of spindle assembly, we attached purified chromosomes to the functionalized and passivated glass coverslip of a microscope flow chamber (Methods). We then added *Xenopus* egg extract supplemented with fluorescent tubulin and EB1 to label microtubules and their growing plus-ends, respectively (Fig. 1a schematic). The action of molecular motors was inhibited using the drug vanadate.

**Figure 1.**
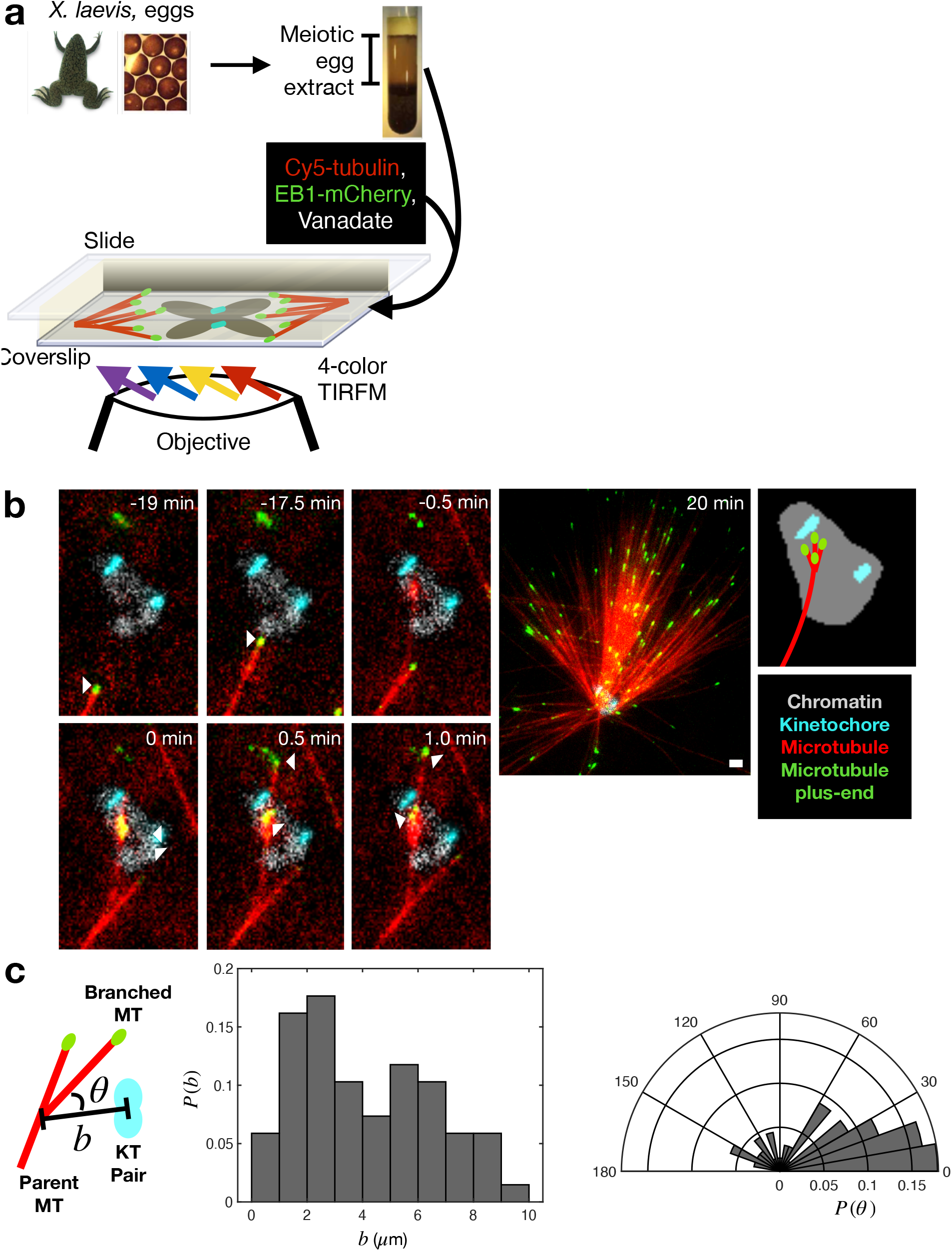
*Ex vivo* reconstitution of branching microtubule nucleation around chromosomes. (a) An illustration of the *ex vivo* reconstitution, which utilizes meiotic cytosol purified from *Xenopus laevis* eggs (Methods). Fluorescent tubulin and EB1 are included to label microtubules and microtubule plus-ends, respectively. Vanadate is used to inhibit motor activity. (b) The initial nucleation events near chromosomes are branches. In the top panel, white markers point to the plus-end of the polymerizing mother microtubule. In the bottom panel, white markers point to initial branching nucleation events. Scale bars are 2 *μ*m. (c) Histogram of distance to the nearest kinetochore pair and polar histogram of angle towards the nearest kinetochore pair for up to the first 10 branching nucleation events around chromosomes. Data are from 11 chromosome clusters across 5 extract preparations. *n* = 68 branching nucleation events.

Using 4-color time-lapse TIRFM, we observed that branched microtubule networks form around chromosomes (Fig. 1b, Movie S1). When the extract solution was first added to surface-bound chromosomes, a very small number of microtubules in the extract nucleated *de novo*, i.e., independent of chromosomes, and were visible at low density throughout the imaging field regardless of whether a chromosome was nearby (Fig. S1a). When a *de novo* microtubule happened to reach a chromosome, a sudden burst of microtubule generation occurred at the chromosome and microtubules appeared to nucleate from it via branching microtubule nucleation (Fig. 1b, Fig. S1b, Movie S1). These chromosomal microtubules formed branched networks of overall uniform polarity (Fig. 1b), similar to the networks observed in previous studies of branching nucleation^25,38^. 50% of these early nucleation events occurred within 4 *μ*m of a kinetochore and 55% of these branched microtubules grew at an angle less than 40° with respect to the kinetochore (Fig. 1c). These results demonstrate that microtubule nucleation for meiotic spindle assembly is initiated near and towards kinetochores.

### Models of branching microtubule nucleation in a uniform field of SAFs

To establish a starting point for our model, we considered the simpler case of branching nucleation uniform field of SAFs. This condition can be probed experimentally by flowing *Xenopus* egg extract supplemented with the non-hydrolyzable Ran mutant RanQ69L into a microscope flow chamber, which creates a uniform field of SAFs for microtubules to branch in (Fig. 2a, Movie S2). We first observe the nucleation of a *de novo* mother microtubule that then seeds the formation of a polar branched network (Movie S2).

**Figure 2.**
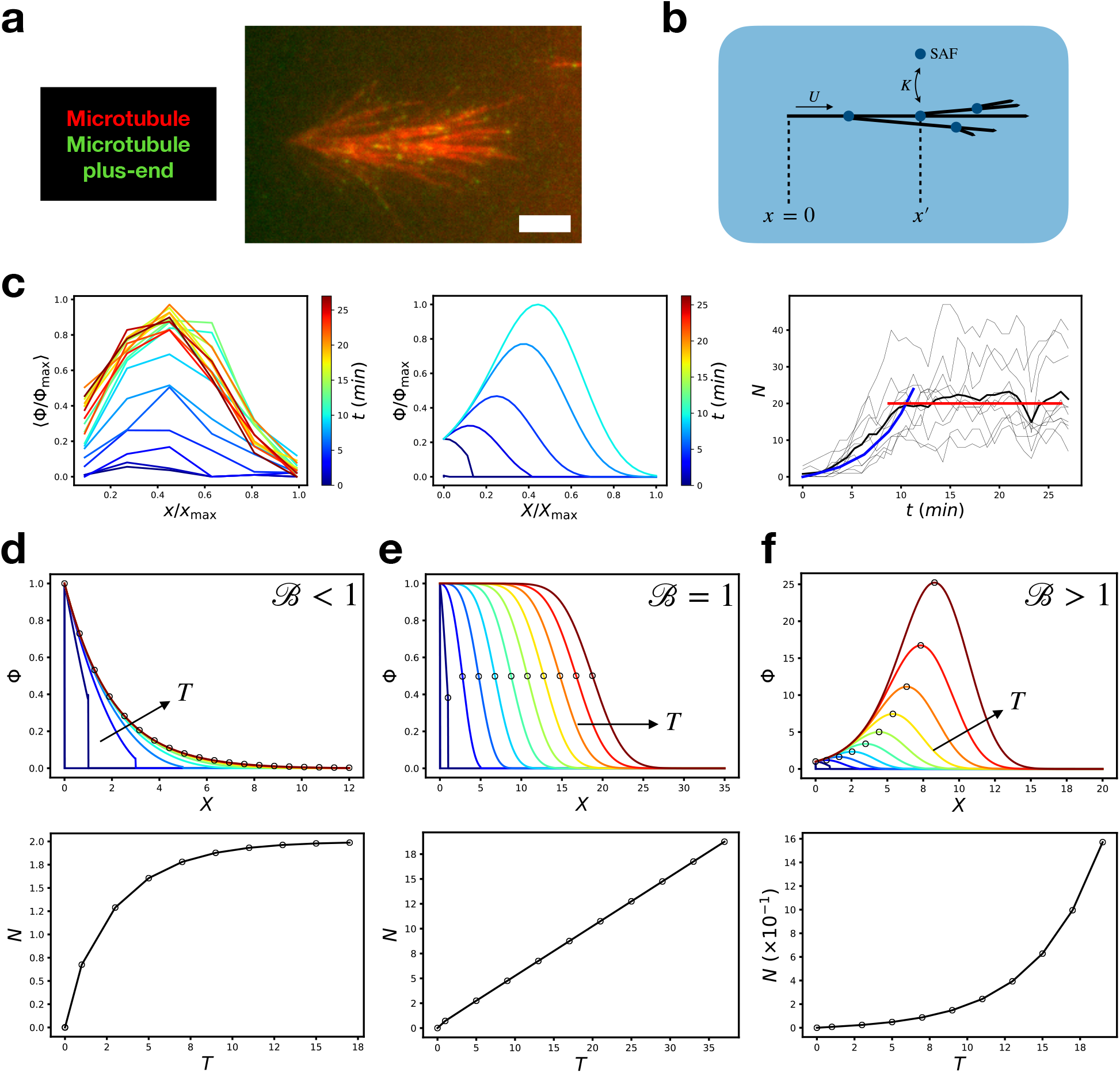
Branching microtubule nucleation in a uniform field of SAFs: experiment and theory. (a) Snapshot of a branched network visualized using TIRFM in a uniform field of SAFs. Scale bar is 5 *μ*m. (b) Schematic of the branching nucleation model. (c) Experimentally measured plus-end distribution (left panel, n = 10 branched networks across 5 different extract preparations) compared with theoretically predicted plus-end distribution using *B* = 1.7 (middle panel). In the right panel, the black curve shows the experimentally measured average number of microtubules, which increases exponentially over the first ~ 10 min before saturating. The blue curve is the theoretical prediction using *B* = 1.7 (*R*^2^ = 0.90), while the red curve shows the average number of microtubules after saturation. (d) For *B* < 1, the plus-end distribution reaches a bounded stationary state. (e) For *B* = 1, the plus-end distribution propagates as a constant density wave. (f) For *B* > 1, the plus-end distribution propagates as an autocatalytic growing front.

To quantify the organization of these branched networks, we measured the distribution of microtubule plus-end positions projected along the axis of the mother microtubule at different times after the addition of extract (Fig. 2b, Methods). We measure the spatial distribution of plus-ends, as opposed to total tubulin density, to distinguish between the nucleation of new microtubules and the polymerization of existing ones. We normalized each plus-end distribution by the maximum length of the final branched network and averaged over *n* = 10 networks to obtain the final result (Fig. 2c left). These branched networks grow autocatalytically for the first ~ 10 min after the addition of extract, consistent with previous work in *Xenopus* egg extract^38^, and then saturate to a stationary state of constant average microtubule number (Fig. 2c right).

To rationalize the structure of these branched networks, we developed a one-dimensional mathematical model of branching nucleation (Supplementary Theory). In the uniform field limit, our model admits an exact solution for the dimensionless plus-end distribution *Φ* as a function of position *X* and time *T*,

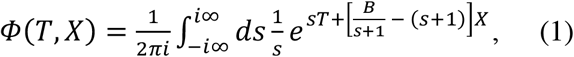

which is parameterized only by the “branching number” *B* = *Kc*_0_*U*/*f_c_*, where *K* is a binding constant of SAFs to microtubules, *c*_0_ is the SAF concentration, *U* is the polymerization speed, and *f_c_* is the catastrophe frequency. Therefore, *B* representations the competition between microtubule nucleation due to branching and microtubule turnover due to catastrophes. The number of microtubules is given by 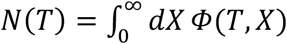.

For *B* < 1, microtubule turnover outcompetes branching nucleation, so microtubule plus-ends can only propagate a finite distance along the mother microtubule (Fig. 2d). After sufficient time, the statistically stationary distribution *Φ*(*T* → ∞, *X*) = *e*^(*B*−1)*X*^ is achieved. When *B* = 1, microtubule turnover balances branching nucleation perfectly, and a branched network of constant microtubule density can propagate indefinitely (Fig. 2e). These constant density waves have been observed for large growing asters in *Xenopus* extracts^39^, where the authors offer a similar physical interpretation and show numerical results. By deriving an explicit formula, we see that their system corresponds to equation (1) when *B* = 1. If *B* > 1, branching microtubule nucleation outcompetes microtubule turnover, and an autocatalytic branched network forms that can propagate indefinitely (Fig. 2f). We find that our experimentally measured plus-end distribution (Fig. 2c left) is well described by our theory with *B* = 1.7 (Fig. 2c middle) during the autocatalytic growth phase, where we used *U* = 8 *μ*m/min^38^ to convert to physical time scales.

After autocatalytic growth, the plus-end distribution reaches a statistically stationary state that is still consistent with *B* > 1 organization. If saturation were due to increasing microtubule turnover, the organization of the plus-end distribution would have to change to that predicted by *B* < 1, which we do not observe experimentally. Therefore, we attribute the reason for network saturation as owing to a limited pool of nucleating factors, and not to increasing microtubule turnover.

### Model of branching microtubule nucleation near chromosomes

To explain how chromosomes might generate branched networks, we modified our theoretical model to include the RanGTP pathway. Chromosomes release RanGTP at a flux *J* into the extract, where it can be hydrolyzed into its inactive RanGDP form at a rate *k_H_*, or it can bind to importin molecules that sequester SAFs, allowing SAFs to promote microtubule branching nucleation (Fig. 3a). This generates a concentration gradient of free SAFs centered at the chromosome. The simplest way to incorporate this pathway into our theoretical framework is to make the change 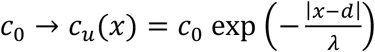, where *λ* is the length scale of the resultant exponential SAF gradient and *d* is the distance between the initial *de novo* nucleation event and the chromosome (Supplementary Theory). In applying our one-dimensional model to the experiments, we are assuming the *de novo* mother microtubule points towards the center of the chromosome, when in reality it is offset by an angle. However, this positive angle is always small in our experiments (14 ± 10°, *n* = 10). Moreover, the gradient biases the branched network to point towards its center, since nucleation events are proportional to the SAF concentration. All the initial *de novo* microtubule needs to do is get close enough, after which the gradient will automatically generate a polar branched network directed at the chromosome. This idea both justifies the use of a one-dimensional model and helps rationalize why the initial branches are directed at chromosomes (Fig. 1c).

**Figure 3.**
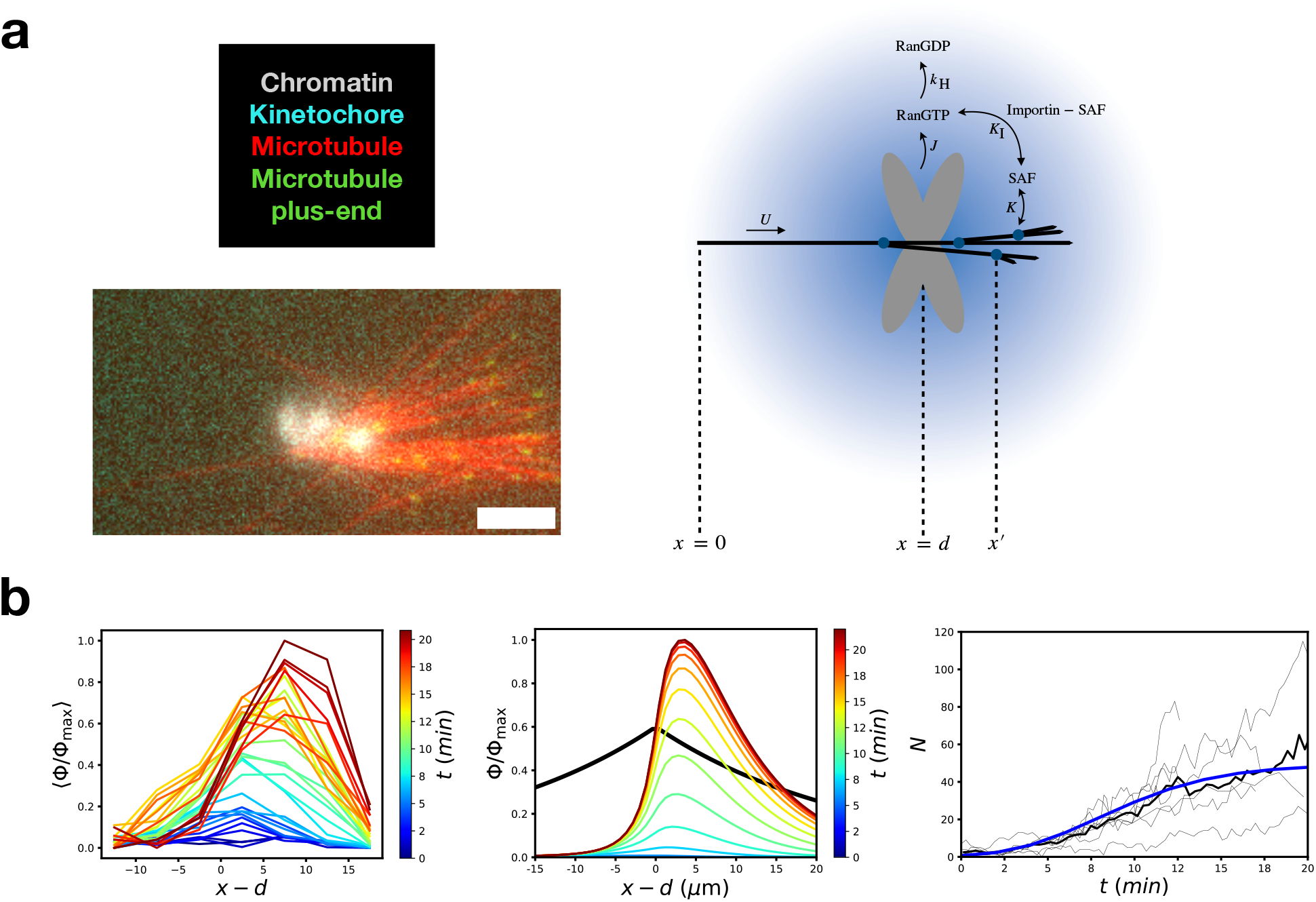
Branching microtubule nucleation near chromosomes: experiment and theory. (a) Model of chromosomal branching microtubule nucleation and representative experimental snapshot. RanGTP is released at chromosomes and can either be hydrolyzed into RanGDP or bind the importin molecules that sequester SAFs, allowing them to promote microtubule branching nucleation. Scale bar is 5 *μ*m. (b) Experimentally measured plus-end distribution (left panel, n = 10 branched networks across 7 different extract preparations) compared with theoretically predicted plus-end distribution using *B* = 2 and *Λ* = 3 (middle panel). The black curve shows the profile of the predicted SAF gradient. In the right panel, the experimentally measured average number of microtubules (black curve) is compared with the theoretical prediction using *B* = 2 and *Λ* = 3 (blue curve, *R*^2^ = 0.95).

In addition to the branching number *B*, we now also have the geometric ratio *Λ* = *λ*/〈*ℓ*〉 as a model parameter. We directly measure *d* from experiments and compute plus-end distributions as a function of *x* − *d*. This model no longer admits a closed-form solution, so we solve numerically for the plus-end distribution. We find that the experimental (Fig. 3b left) and theoretical (Fig. 3b middle) distributions agree qualitatively with each other with *B* = 2 and *Λ* = 3, where we used 〈*ℓ*〉 = 8 *μ*m^27^ and *U* = 8 *μ*m/min^38^ to convert to physical length and time scales. Thus we estimate a SAF gradient length scale of *λ* ≈ 24 *μ*m, which agrees with previous estimates in *Xenopus* systems^27^. We see that a telltale sign of branching nucleation in a SAF gradient is that the plus-end distribution peaks downstream of the chromosome; there is comparatively little plus-end density upstream of the chromosome. Because both our experiments and theory have this feature, we attribute this asymmetry of the plus-end distribution to the spatial symmetry breaking caused by the SAF gradient. Indeed, in a uniform field of SAFs the plus-end distributions are roughly symmetric (Fig. 2c). Moreover, because the SAF gradient is finite in extent, as opposed to the uniform field model, there is now a limiting pool of nucleating factors bounded by 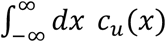, and thus our theory naturally captures both the early time amplification and late time saturation of the total number of microtubules versus time (Fig. 3b right).

Thus, we conclude that the observed organization of chromosomal microtubules in our system is consistent with a model of branching microtubule nucleation in an effector gradient spatially regulated by chromosomes.

### Branching microtubule nucleation is the principal source of chromosomal microtubules

Is branching microtubule nucleation indeed the principal source of chromosomal microtubule nucleation? In order to assess this question, we individually immunodepleted the essential branching factors TPX2 and augmin from the extract^25^ and performed the TIRFM-based assay of acentrosomal spindle assembly from chromosomes (Fig. 4). In control experiments, using a random IgG antibody for immunodepletion, 34 ± 26 % of chromosomes generated branched networks (mean ± standard deviation, *n* = 27 fields, totaling 213 chromosome clusters) (Figs. 4a top, S2 left). In contrast, we rarely saw microtubule networks generated at chromosomes after depletion of either TPX2 or augmin. Less than 1 ± 2 % of chromosomes for either augmin depletion or for TPX2 depletion showed branching (Figs. 4a middle and bottom, S2 right, S3, Movies S3, S4) (mean ± standard deviation, *n* > 10 fields per condition, with > 135 chromosome clusters per condition). These results suggest that branching microtubule nucleation, mediated by augmin and TPX2 in *Xenopus* egg extract, is the key pathway to generate microtubules from chromosomes. Therefore, in meiotic *Xenopus* spindles, chromosomal microtubule nucleation and branching microtubule nucleation are one and the same, and represent the molecular origin of spindle microtubules.

**Figure 4.**
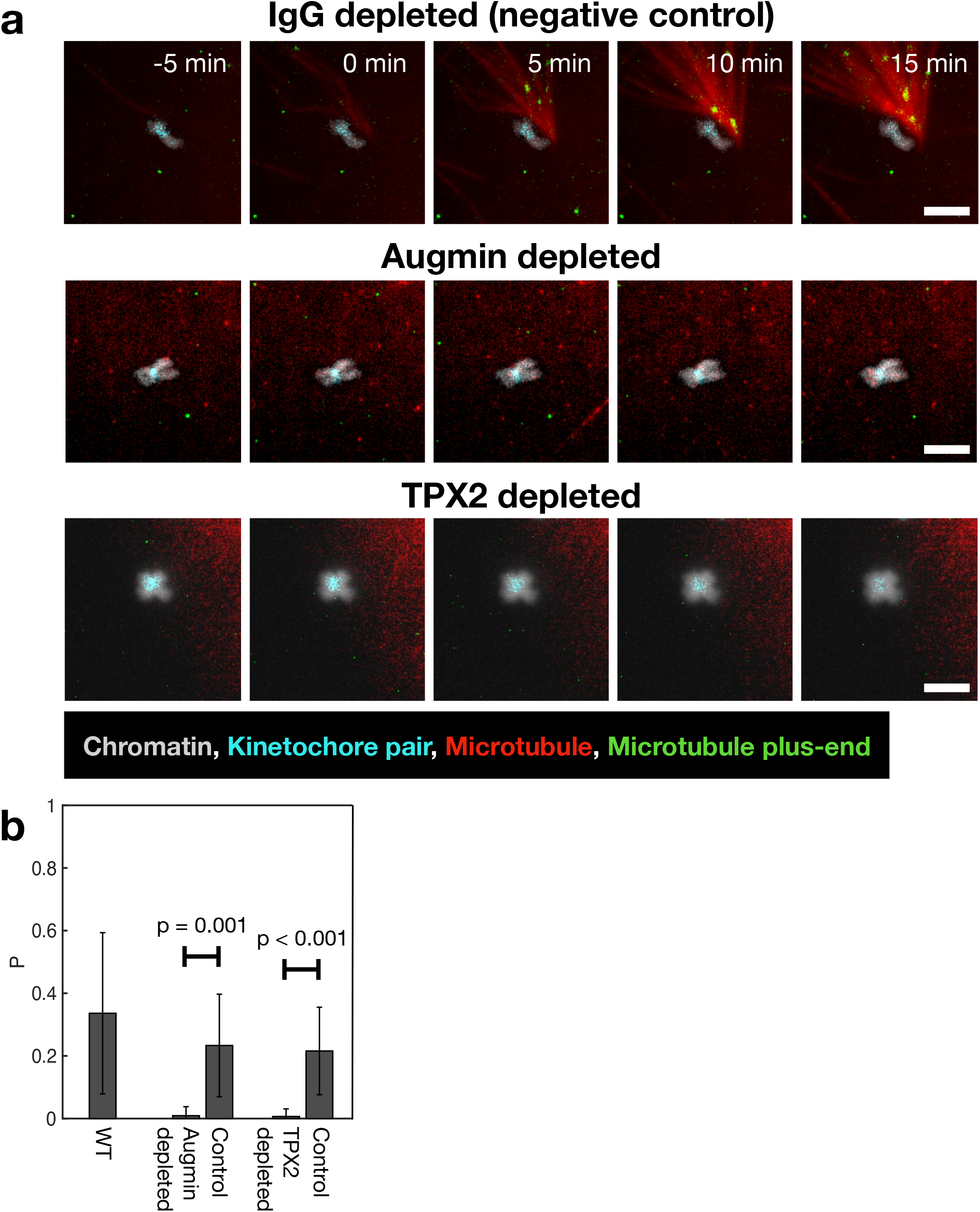
Branching microtubule nucleation mediated by TPX2 and augmin is the chief source of chromosomal microtubule networks. (a) Chromosomes in each of the three conditions tested: IgG depleted, augmin depleted, and TPX2 depleted, visualized using TIRFM. Chromosomal microtubule networks were not generated when augmin or TPX2 was depleted. Scale bars are 5 *μ*m. (b) Fraction of chromosomes that form microtubule networks in each condition. Error bars are standard deviations across multiple fields. *n* > 10 fields and *n* > 100 chromosomes per condition. Data are from 3 to 4 extract preparations per condition. P-values reported are from the two-sample Kolmogorov-Smirnov test. Quantification of average microtubule mass over time per condition is in Fig. S3.

### Reconstituting acentrosomal spindle assembly and organization in Xenopus egg extract

Having determined where, when, and how microtubules are made at chromosomes in the absence of motor activity, we sought to see what happens when motor activity is not inhibited in our assay. We observed that the initial microtubule nucleation events are still branches that grow near and towards kinetochores (Fig. 5a left, Movie S5), but that motor activity then reorganizes these branched networks to produce multipolar microtubule networks, i.e., spindles (Fig. 5a right, Movie S5). The spindles that assembled in the presence of motor activity (Fig. 5b) stand in contrast to the polar branched networks seen in Figs. 1–3. The majority of spindles were bipolar (Fig. 5c), similar to spindles generated from chromatin in *Xenopus* egg extract^6^. The emergent bipolar structure is particularly striking when the spindle is allowed to form around chromosomes in the three-dimensional bulk extract, which we captured using epifluorescence microscopy (Fig. 5d). These results demonstrate that chromosomes alone can assemble acentrosomal spindles via branching microtubule nucleation and motor activity (Fig. 5e).

**Figure 5.**
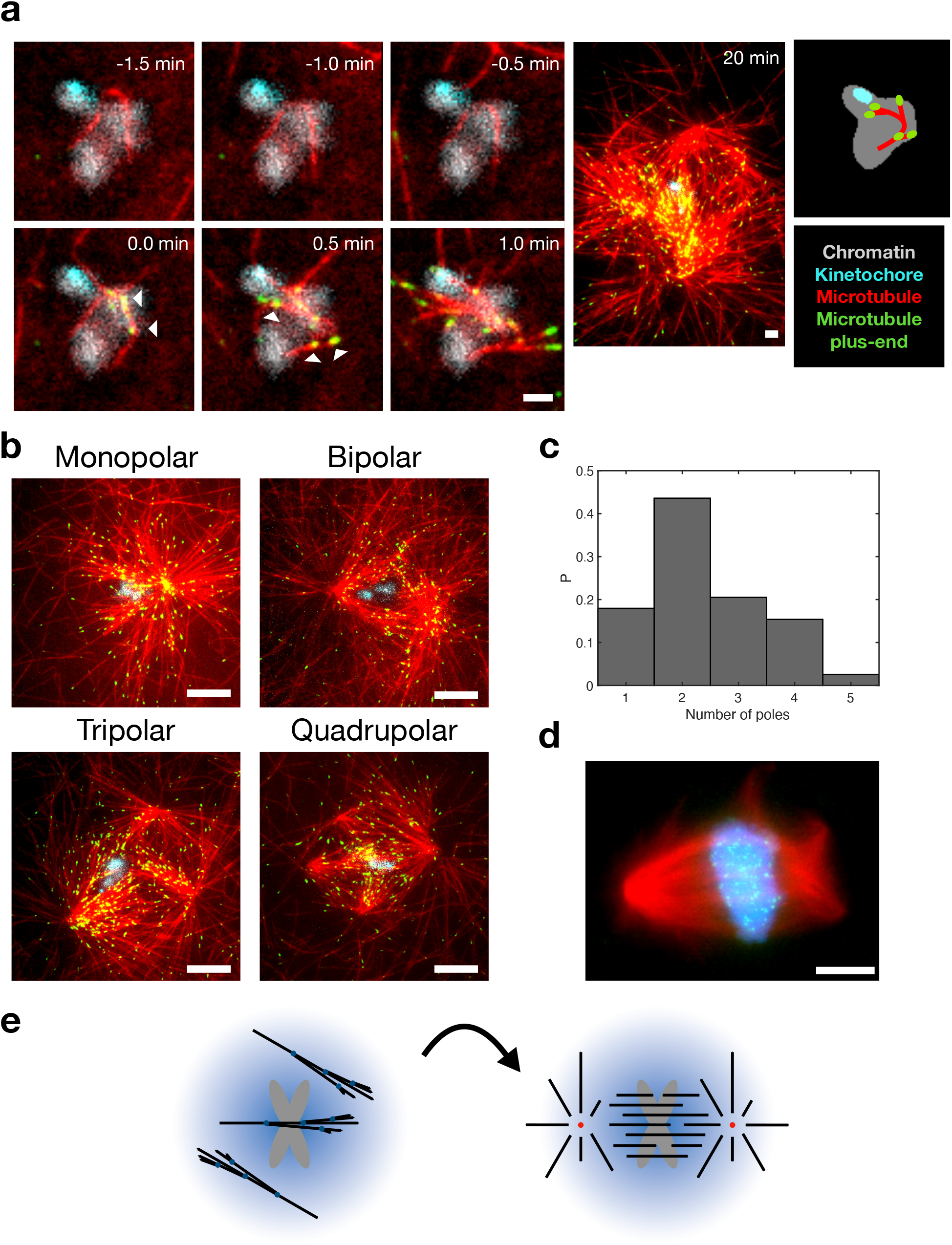
In the presence of motor activity, chromosomes generate polar spindles. (a) Branching microtubule nucleation leads to the formation of branched networks at chromosomes, which are eventually organized into a spindle by motor activity. Time 0 is the first appearance of a branch. White markers point to initial branching nucleation events. Scale bars are 2 *μ*m. (b) TIRFM visualization of mono-, bi-, tri-, and quadrupolar spindles assembled around purified chromosome clusters. Scale bars are 10 *μ*m. (c) Histogram showing distribution of number of poles for chromosomal microtubule networks. *n* = 39 spindles. (d) Fixed epifluorescence image of a three-dimensional bipolar spindle generated around purified chromosomes. Scale bar is 10 *μ*m. (e) Schematic of our general model for acentrosomal spindle assembly. First, *de novo* microtubules find their way into the SAF gradient generated by chromosomes. Branching microtubule nucleation occurs along these first mother microtubules, generating branched networks. These branched networks are then self-organized by molecular motors into a bipolar spindle.

We went further with this assay and studied the dynamics of spindle formation around chromosomes (Fig. 6a, Movie S6). By measuring the total tubulin intensity and number of EB1 spots over time in a 40 *μ*m x 40 *μ*m box around chromosomes, we found that the total microtubule mass and number of microtubules in these spindles plateau ~ 10 min after the onset of spindle assembly (Fig. 6b, 6c), consistent with our experiments using vanadate to inhibit motors (Figs. 2c right, 3b right). The time to plateau is also consistent with previous findings based on similar experiments with chromatin beads^36,40^. Because both the number of microtubules and total microtubule mass plateau around the same time (Fig. 6b, 6c), this further supports the idea that microtubule mass in the spindle is limited by available nucleators, and not by tubulin availability or changing microtubule polymerization dynamics.

**Figure 6.**
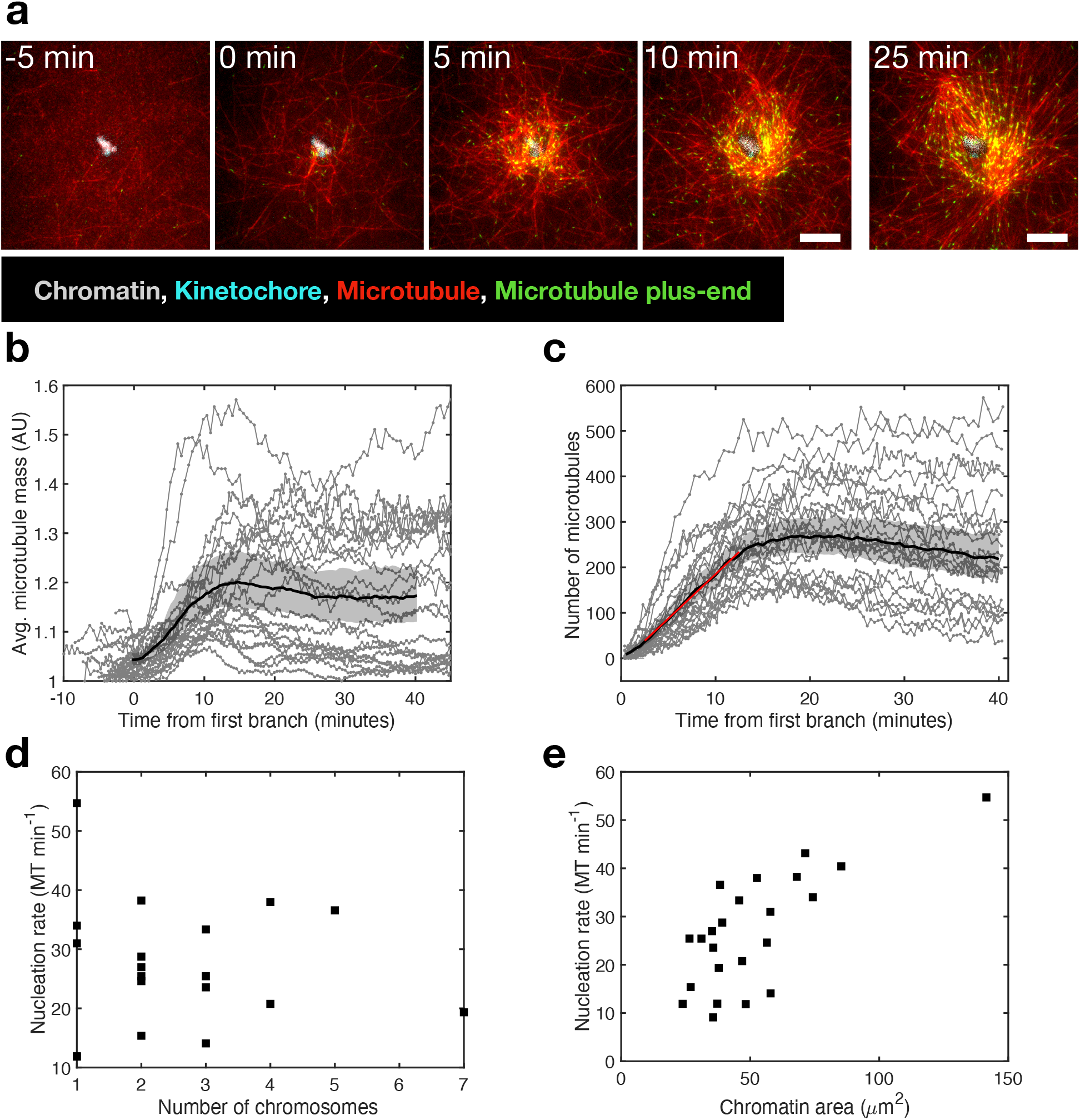
Kinetics of microtubule nucleation during spindle assembly around chromosomes. (a) Timelapse TIRFM of a bipolar spindle assembling around chromosomes. Scale bar is 10 *μ*m. (b) Microtubule mass and (c) number of microtubules versus time during spindle assembly. Microtubule mass and number plateau at ~ 10 min. *n* = 23 spindles across 7 extract preparations. Shaded regions represent 95% bootstrap confidence intervals. In (c), the red curve is a linear fit, giving an effective microtubule nucleation rate of *k* = 19.5 ± 0.54 microtubules/min (mean ± 95% confidence bounds) (*R*^2^ > 0.99) (d) Nucleation rate in the spindle does not significantly correlate with the number of visible kinetochore pairs in the chromosome cluster. Pearson correlation coefficient = −0.07, p = 0.78. *n* = 19 spindles across 7 extract preparations. (e) The nucleation rate in the spindle correlates with the two-dimensional projected area of the chromatin in the chromosome cluster. Pearson correlation coefficient = 0.73, p < 0.001. *n* = 23 spindles across 7 extract preparations.

We next asked if acentrosomal spindle assembly relies on the number of chromosomes or the amount of chromatin. To determine the number of chromosomes, we counted the number of kinetochore pairs, although we note that there could be more kinetochores above the focal plane of TIRFM. We found that there was no significant correlation between nucleation rate and the number of chromosomes, which we counted in each chromosome cluster (Fig. 6d; Pearson correlation coefficient = −0.07, p = 0.78). In contrast, there was a positive and significant correlation between the nucleation rate and the two-dimensional projected area of the chromatin in each chromosome cluster (Fig. 6e; Pearson correlation coefficient = 0.73, p < 0.001). Thus, these results show that the amount of chromatin in chromosomes sets the nucleation rate of microtubules in the spindle, independent of the number of chromosomes. This is consistent with our theoretical model, since the branching number *B* scales with the SAF concentration *c*_0_, which in turn scales with the flux of RanGTP *J* produced by chromosomes. Since *J* is proportional to the total surface area of chromatin, we expect the nucleation rate to be proportional to the total chromatin area.

Lastly, we investigated whether microtubules polymerized directly through chromosomes. To test this idea, we let acentrosomal spindles form around chromosomes for 20 min and then plotted the mean intensity of the tubulin channel and all microtubule tracks around chromosomes (Fig. S4. Interestingly, the microtubule tracks suggest that microtubules can pass through less compact regions of chromosomes, such as the chromosome arms. In stark contrast, the tubulin intensity and microtubule tracks both featured voids at the kinetochores. No microtubules out of thousands in a spindle passed through the kinetochores of a chromosome—even the tracks that go toward the kinetochores. These results suggest that microtubules cannot polymerize through kinetochores.

## DISCUSSION

In this work, we made several experimental and theoretical advances to investigate acentrosomal spindle assembly, and can now propose the following model. First, *de novo* microtubules nucleate randomly throughout the cytoplasm. When one of these microtubules finds its way near chromosomes, branched microtubules nucleate from it due to the action of the SAF gradient around chromosomes. In the absence of motors, branched microtubule networks of uniform polarity directed at kinetochores form (Fig. 3). When motors are present, self-organized multipolar microtubule networks form around chromosomes, with the most probable organization being a bipolar spindle (Fig. 5).

Previous work on meiotic spindle assembly suggested that early microtubules formed at random locations and orientations, based on imaging techniques that could not resolve individual filaments^6,36,41^. In contrast, because TIRFM provides excellent signal-to-noise ratio, we see that single *de novo* microtubules seed the formation of spindles at chromosomes. These microtubules generate many more branched microtubules that are no longer randomly oriented, but nucleate near and are oriented towards kinetochores, which we attribute to the spatial nucleation bias generated by the RanGTP-induced SAF gradient. This gives rise to two conclusions. First, we show that microtubule generation around chromosomes during meiosis is attributable to RanGTP-mediated branching microtubule nucleation. Second, because branching microtubule nucleation in a SAF gradient begins near and towards kinetochores, the capture of chromosomes by the spindle during meiosis is made more feasible. In fact, how kinetochores are captured and kinetochore fibers form in the absence of centrosomes is an open question. Our results, for the first time, show how this is possible: with polarized, branched networks that point toward kinetochores due to a spatially biasing SAF gradient. We posit that this same mechanism occurs in centrosomal spindles, where it would serve to complement the classical search and capture model of centrosomal microtubules by having microtubules form via branching nucleation directly from chromosomes.

We note that previous work in mammalian eggs showed that small microtubule organizing centers (MTOCs) consisting of many microtubules seeded the subsequent assembly of the spindle after breakdown of the nuclear envelope in a Ran-dependent manner^42^. Whether branching microtubule nucleation is also significant for these small MTOCs, and whether a RanGTP gradient is involved, remains to be determined.

Because we used purified chromosomes, we were able to observe that no microtubule tracks pass through the kinetochores in the center of chromosomes, yet tracks do appear to pass through the peripheral chromatin arms (Fig. S4). This finding suggests that microtubules, which are 25 nm in diameter, may pass through gaps within chromatin. It is known that microtubules interact with other proteins on the surface of chromosomes besides kinetochores, such as the chromokinesin motor to generate polar ejection forces that move chromosomes away from the spindle poles^43–47^. In light of our data, the possibility that microtubules interact with proteins or DNA within chromosomes suggests new avenues for investigation into the intersection of chromatin and cytoskeletal biology. Interestingly, it has been recently suggested that chromosomes can regulate their structure to prevent or enhance microtubules from perforating them^48^.

Based on this framework and mechanism we provide here for how microtubule nucleation around chromosomes occurs, the time is now ripe to further investigate how molecular motors organize these initial chromosomal branched networks into a functioning bipolar spindle. This issue is only now starting to be quantitatively addressed^49^.

## ACKNOWLEDGEMENTS

We thank Jake DeLuca, Keith DeLuca, and Jeanne Mick for the CENPA-GFP HeLa cell line, Kayoko Hayashihara and Kiichi Fukui for helpful discussions regarding chromosome isolation, Nachama Sterm for assisting with chromosome counting in purified fractions, Gary Laevsky and the Confocal Imaging Facility, and all members of the Petry, Shaevitz, and Stone labs for advice and input. S.U.S. was supported by NIH NCI NRSA 1F31CA236160 and NHGRI training grant 5T32HG003284. B.G. was funded by the PD Soros fellowship and the NSF GRFP DGE-2039656. M.R.K. was supported by NIGMS training grant T32GM007388. This work was funded by NIH NIA 1DP2GM123493, Pew Scholars Program 00027340, Packard Foundation 201440376, and CPBF NSF PHY-1734030.

## AUTHOR CONTRIBUTIONS

S.U.S. purified chromosomes, designed and performed experiments, and analyzed experimental data. B.G. performed theoretical analysis with assistance from H.A.S., numerical calculations, and analyzed experimental data. M.R.K. assisted in testing chromosome isolation protocols. S.U.S. and B.G. wrote the manuscript. H.A.S, J.W.S., and S.P. contributed to research design, mentoring, and writing the manuscript.

## ETHICS

Animal care was done in accordance with recommendations in the Guide for the Care and Use of Laboratory Animals of the NIH and the approved Institutional Animal Care and Use Committee (IACUC) protocol 1941-16 of Princeton University. Data and code used are available upon request.

## SUPPLEMENTAL FIGURES AND LEGENDS

**Figure S1.**
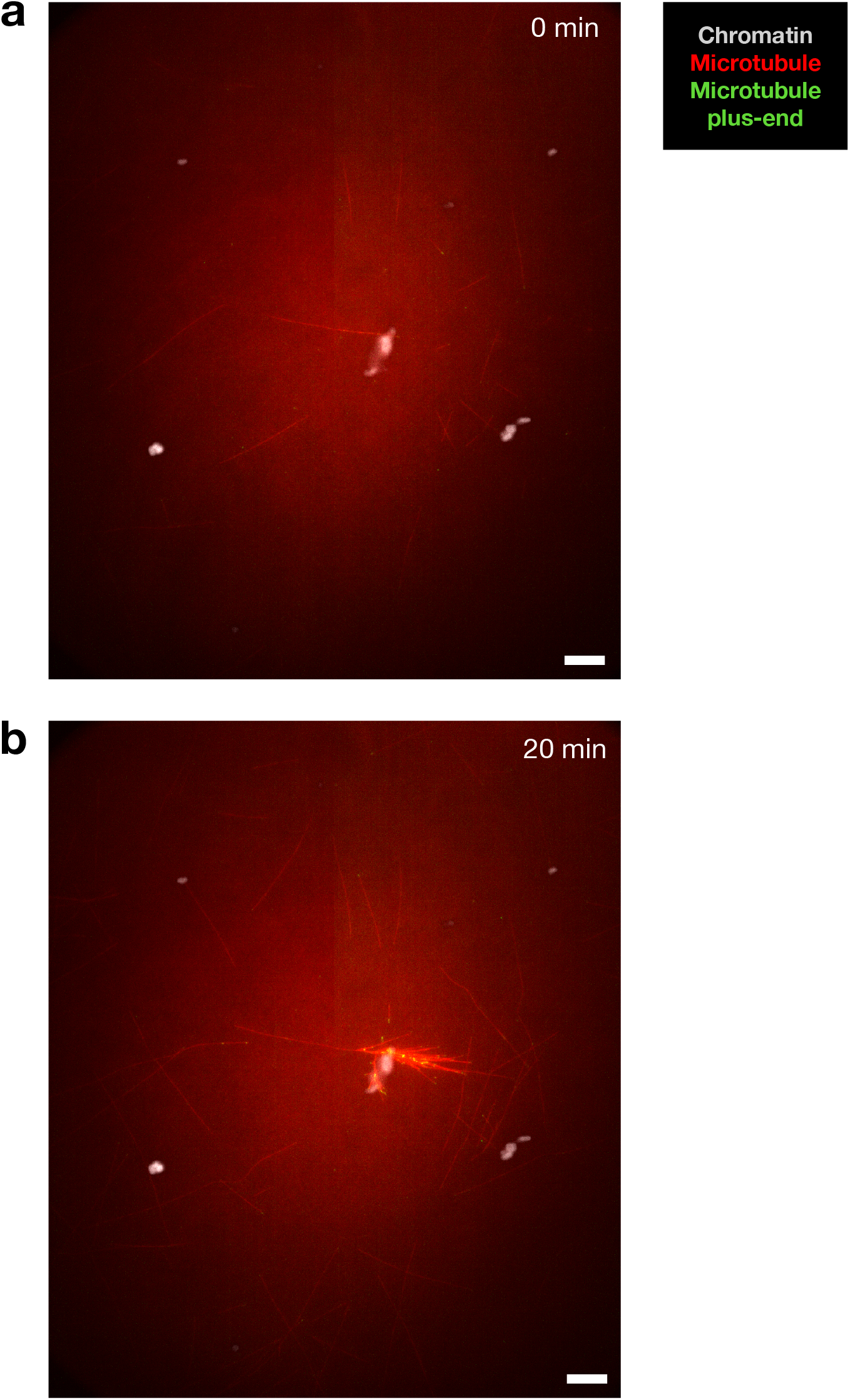
Starter *de novo* microtubules randomly nucleate to initiate branched networks near chromosomes. Scale bars are 10 *μ*m.

**Figure S2.**
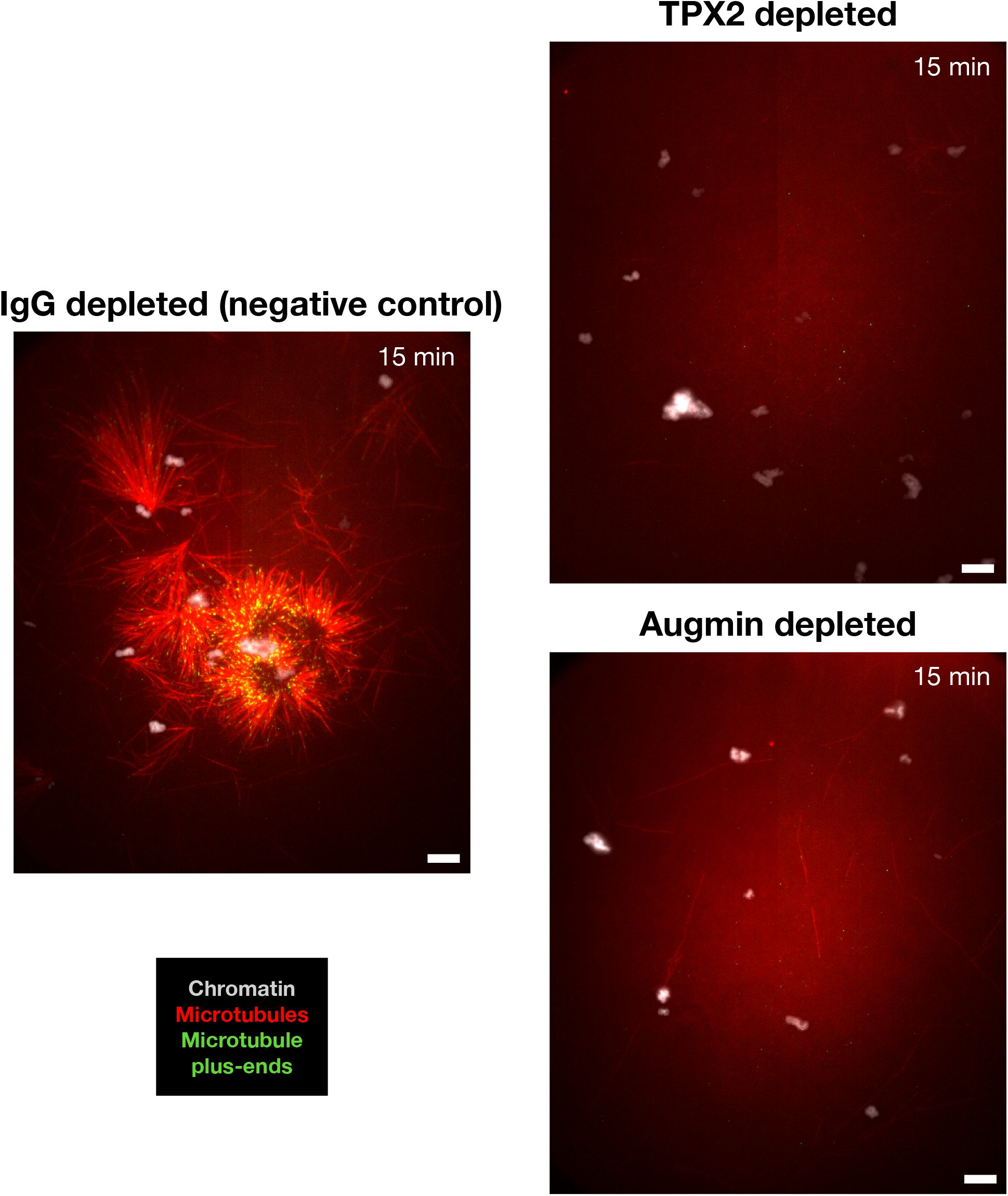
Entire field of view of TPX2 and augmin depletion experiments. Scale bars are 10 *μ*m.

**Figure S3.**
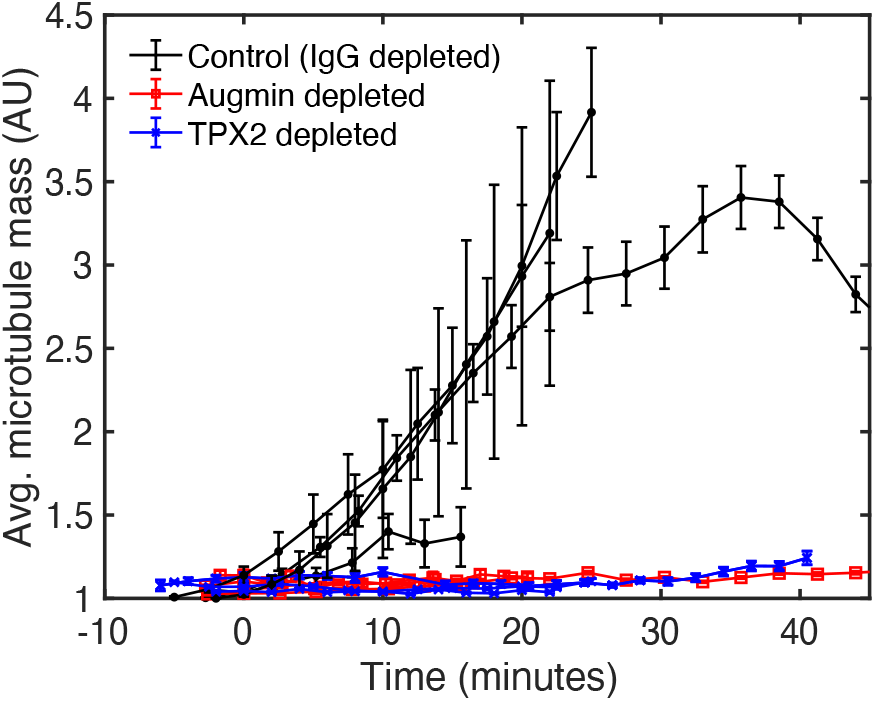
Average microtubule mass over time for chromosomal microtubule networks in control depleted, augmin depleted, and TPX2 depleted extracts. Each curve represents a separate extract experiment. Error bars are taken across chromosomes and are standard error of the mean. Chromosomes that generated networks in the control conditions were included and all chromosomes in the depletion conditions were included. Only one chromosome cluster showed microtubule network formation in either depletion condition (< 1%, see Fig. 4b) out of *n* > 100 chromosomes in each and across all extracts. *n* = 19.7 ± 19.8 chromosomes per curve (mean ± standard deviation).

**Figure S4.**
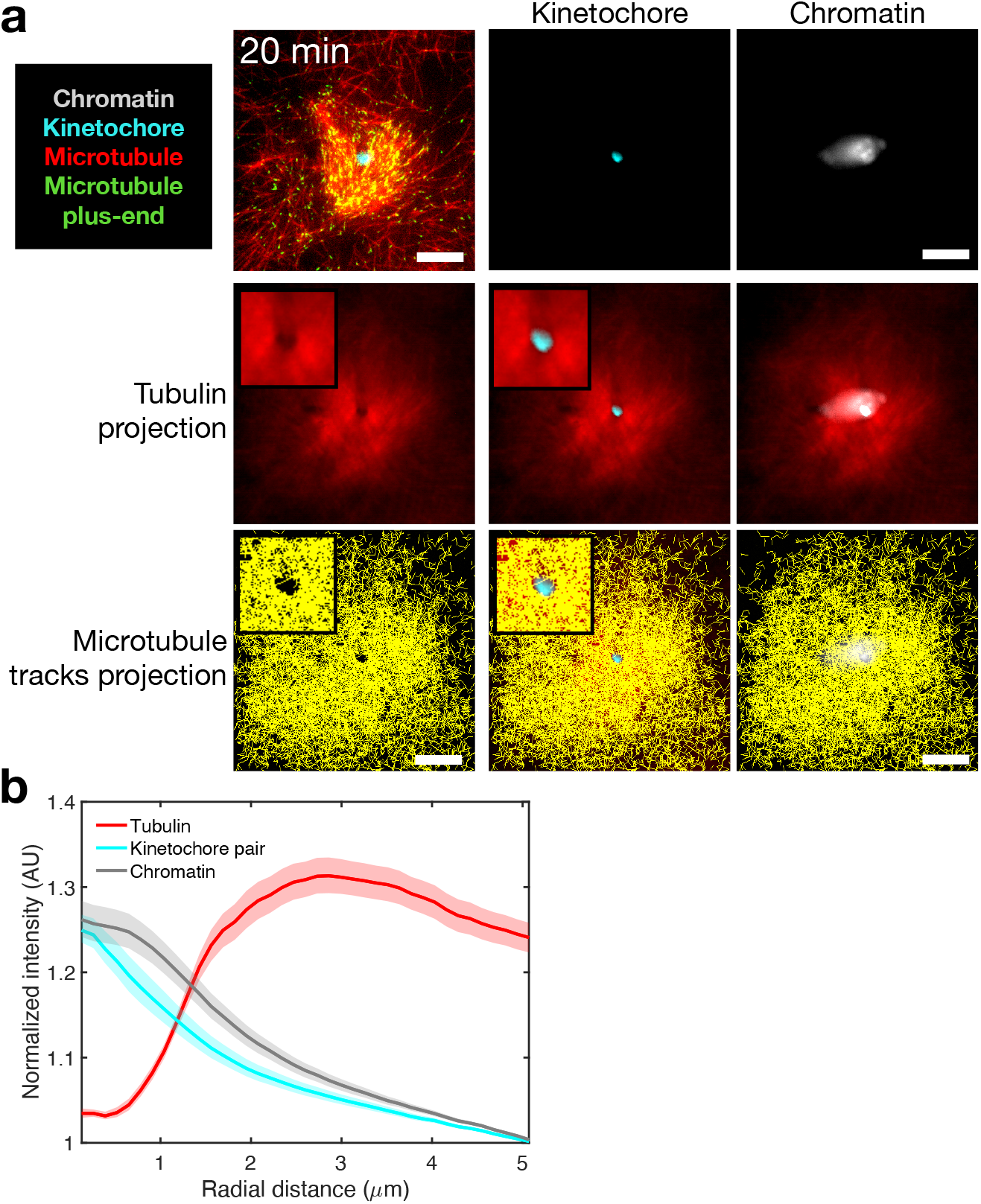
Kinetochores act as a void for microtubule nucleation and polymerization. (a) Images of tubulin mean time projection, kinetochore, chromatin, and microtubule tracks time projection. The bottom right panels show merged images. Both the tubulin mean time and microtubule tracks projections exhibit a void at the kinetochore, because microtubules do not polymerize through the kinetochore. Scale bars are 10 *μ*m. Zoomed-in insets of the void region are 10 *μ*m x 10 *μ*m. (b) Quantification of normalized intensity for kinetochore, tubulin, and chromatin channels. Data are from the 3 chromosome clusters across 2 extracts that had only 1 void visible. Every chromosome cluster featured at least 1 void. Shaded regions represent 95% bootstrap confidence intervals.

## MOVIE LEGENDS

**Movie S1. Branching microtubule nucleation at kinetochores with vanadate.** The field is 30 *μ*m x 30 *μ*m.

**Movie S2. Branching microtubule nucleation in a uniform field of SAFs.** The field is 34 *μ*m x 22 *μ*m.

**Movie S3. Branched microtubule network formation in augmin depleted (left) and control depleted (right) extracts**. The fields are 140 *μ*m x 166 *μ*m.

**Movie S4. Branched microtubule network formation in TPX2 depleted (left) and control depleted (right) extracts**. The fields are 140 *μ*m x 166 *μ*m.

**Movie S5. Branching microtubule nucleation at kinetochores with motor activity.** The field is 30 *μ*m x 30 *μ*m.

**Movie S6. Bipolar spindle assembly around chromosomes.** The field is 40 *μ*m x 40 *μ*m.

## MATERIALS AND METHODS

### Protein expression and purification

EB1-mCherry was purified as previously described^50^. Protein was expressed in E. coli (strain Rosetta 2) for 4 hours at 37 C. Cells were lysed via a french press using an EmulsiFlex (Avestin) in lysis buffer (50 mM NaPO_4_, pH 7.4, 500 mM NaCl, 20 mM imidazole, 2.5 mM PMSF, 6 mM CME, 1 cOmplete™ EDTA-free Protease Inhibitor (Sigma), 1000 U DNAse 1 (Sigma)). Protein was affinity purified from lysate using a HisTrap HP 5 mL column (GE Healthcare) in binding buffer (50 mM NaPO_4_, pH 7.4, 500 mM NaCl, 20 mM imidazole, 2.5 mM PMSF, 6 mM BME). Protein was then eluted using elution buffer (50 mM NaPO_4_, pH 7.4, 500 mM NaCl, 500 mM imidazole, 2.5 mM PMSF, 6 mM BME). Next, peak fractions were pooled and loaded onto a Superdex 200 pg 16/600 gel filtration column, and gel filtration was done in CSF-XB (10 mM HEPES, pH 7.7, 1 mM MgCL2, 100 mM KCl, 5mM EGTA) with 10% (w/v) sucrose.

RanQ69L, used to test the quality of meiotic *Xenopus* egg extract prior to experiments with chromosomes, was also purified as previously described^50^. RanQ69L, tagged on its N-terminus with BFP to improve solubility, was expressed and then lysed as above in lysis buffer (100 mM tris-HCl, pH 8.0, 450 mM NaCl, 1 mM MgCl_2_, 1 mM EDTA, 0.5 mM PMSF, 6 mM BME, 200 *μ*M GTP, 1 cOmplete™ EDTA-free Protease Inhibitor, 1000 U DNAse 1). Protein was then affinity purified from lysate using a StrepTrap HP 5 mL column (GE Healthcare) in binding buffer (100 mM tris-HCl, pH 8.0, 450 mM NaCl, 1 mM MgCl_2_, 1 mM EDTA, 0.5 mM PMSF, 6 mM BME, 200 *μ*M GTP). Bound protein was eluted using elution buffer (100 mM tris-HCl, pH 8.0, 450 mM NaCl, 1 mM MgCl_2_, 1 mM EDTA, 0.5 mM PMSF, 6 mM BME, 200 *μ*M GTP, 2.5 mM D-desthiobiotin). Finally, eluted protein was dialyzed into CSF-XB (10 mM HEPES, pH 7.7, 1 mM MgCL2, 100 mM KCl, 5mM EGTA) with 10% (w/v) sucrose overnight.

Tubulin from bovine brain (PurSolutions) was labeled with Cy5 NHS ester (GE Healthcare) as previously described^51^.

Protein concentration was assessed using SDS-PAGE followed by Coomassie staining against a standard of BSA with known concentrations, or via Bradford dye (Bio-Rad).

### Chromosome isolation

Chromosomes were isolated from mitotic HeLa cells following previous approaches^52–55^. First, CENPA-GFP HeLa cells were synchronized to mitosis via a single or double thymidine block^56^. Prior to collecting cells, cytochalasin D was added to media at a final concentration of 10 mg/mL. Next, cells were collected and swelled in a hypotonic solution of 0.075 M KCl for 20 minutes at 37 C. After this, all work was done at 4 C. The cells were centrifuged at 780 g for 15 minutes. The supernatant as removed and then 25 mL of polyamine solution (15 mM tris-HCl, pH 7.4, 2 mM EDTA, 80 mM KCl, 20 mM NaCl, 0.2 mM spermine, 0.5 mM spermidine, 0.05% (v/v) Empigen BB (Sigma), 7 mM BME, 1 cOmplete™ EDTA-free Protease Inhibitor) was layered over the pellet. The pellet and polyamine solution were kept on ice for 5 minutes and then gently resuspended. Cells were then gently lysed by a Dounce homogenizer (B pestle, 10 passes). The lysate was gently centrifuged at 190 g for 3 minutes, 7/10^th^ of the supernatant taken, and then the supernatant was more strongly centrifuged at 1750 g for 20 minutes onto a 70% (v/v) glycerol cushion of the polyamine solution. The layer of chromosomes above the cushion was gently resuspended with the cushion. Next, the resuspended cushion sample and 30 mL Percoll buffer (5 mM tris-HCl, pH 7.4, 20 mM EDTA, 20mM KCl, 0.8 mM spermine, 2.25 mM spermidine, 1% (v/v) thiodiethanol, 0.05% (v/v) Empigen BB, 89% (v/v) Percoll (GE healthcare), 1 cOmplete™ EDTA-free Protease Inhibitor) were added to a Dounce homogenizer to a final volume of about 35 mL and gently homogenized (B pestle, 10 passes). Then, the homogenized solution was brought to 55 mL with more Percoll buffer and centrifuged at 48400 g for 30 minutes in a 45 Ti rotor (Beckman). Chromosomes appeared as a faint band about 1/5^th^ from the bottom of the centrifuge tube. The Percoll gradient was manually fractionated and fractions were imaged for chromosomes via epifluorescence. Chromosome-rich fractions were pooled, diluted three-fold in dilution buffer (5 mM tris-HCl, pH 7.4, 20 mM EDTA, 20mM KCl, 0.8 mM spermine, 2.25 mM spermidine, 1% (v/v) thiodiethanol, 0.05% Empigen BB, 1 cOmplete™ EDTA-free Protease Inhibitor), and centrifuged at 1250 g for 20 minutes onto a 2M sucrose CSF-XB (10 mM HEPES, pH 7.7, 1 mM MgCL2, 100 mM KCl, 5mM EGTA, 2M sucrose) cushion twice, resuspending the first cushion and sample with CSF-XB (10 mM HEPES, pH 7.7, 1 mM MgCL2, 100 mM KCl, 5mM EGTA). The final cushion and sample were gently resuspended and then aliquoted and flash frozen.

### Surface chemistry for chromosome attachment

Flow chambers were made using double-sided tape to create a rectangular chamber between a glass slide and a coverslip. Anti-ds DNA antibody (Abcam, ab27156) was flowed in at 0.1 to 0.17 mg/mL and allowed to adhere to coverslips for 10 minutes. Excess antibody was washed out three times using CSF-XB (10 mM HEPES, pH 7.7, 1 mM MgCL2, 100 mM KCl, 5mM EGTA) with 10% (w/v) sucrose, and then the coverslip surface was passivated using *κ*-casein at 1 mg/mL, incubated for 10 minutes. Next, diluted, purified chromosomes stained with DAPI (0.1-1 *μ*g/mL) were flowed into the chamber and allowed to bind for 10 to 20 minutes. Unbound chromosomes were washed out three times using CSF-XB (10 mM HEPES, pH 7.7, 1 mM MgCL2, 100 mM KCl, 5mM EGTA) with 10% (w/v) sucrose. Meiotic *Xenopus* egg extract was flowed into the chamber and chromosomal microtubule nucleation visualized using TIRFM.

Coverslips (no. 1.5, Fisher or equivalent) were silanized for immunodepletion and drug inhibition experiments following an existing protocol^51^. Briefly, coverslips were sonicated using a bath sonicator for 5 minutes in 1 M NaOH, rinsed with DI water twice, sonicated in DI water for 5 minutes, and then dried using inert nitrogen gas. Coverslips were then silanized for 1 hour using a 0.05% (v/v) dichloro(dimethyl)silane (DDS) solution in trichloroethylene (TCE), with gentle stirring. Coverslips were then sonicated through a series of methanol baths for 5, 15, and 30 minutes, and finally dried using inert nitrogen gas. Silanized coverslips were stored in clean, sealed glass containers and used within one week.

### Meiotic *Xenopus laevis* egg extract purification, protein immunodepletion, and drug inhibition

Meiotic *Xenopus laevis* egg extract, also known as M-phase, metaphase arrested, or CSF extract, was prepared from *Xenopus laevis* eggs according to previously described protocols^57,58^. Egg extract was diluted no more than 75% for all TIRFM experiments, and prepared as 20 *μ*L reactions containing 15 *μ*L extract, 0.89 *μ*M Cy5-labeled tubulin, and 200 *μ*M EB1-mCherry. CSF-XB (10 mM HEPES, pH 7.7, 1 mM MgCL2, 100 mM KCl, 5mM EGTA) with 10% (w/v) sucrose was added to bring final dilution down to 75%, as needed. Freshly prepared extract was tested for its ability to generate branched microtubule networks before further experiments were done or immunodepletion was started by visually comparing reactions without and with 10 *μ*M BFP-RanQ69L. For depletion of TPX2 or augmin from egg extract, 72 *μ*g of purified anti-TPX2 or anti-Haus1 antibody, as used previously^38^, was coupled overnight to 300 *μ*L protein A magnetic beads (Dynabeads, ThermoFisher). The following day, 150 *μ*L fresh egg extract was depleted of either target protein in three rounds of washes with beads, using 100 *μ*L of beads per wash, 20 minutes per round. Control depletions were done using rabbit IgG antibody. The efficiency of depletion using these antibodies was previously determined by western blot and confirmed by assaying for Ran aster generation without and with 10 *μ*M BFP-RanQ69L. Protein and control depleted extracts were imaged simultaneously using multichannel flow chambers made using multiple pieces of double-sided tape.

For inhibiting aurora B kinase, AZD1152^59^ (Sigma) was added to fresh egg extract at a final concentration of 1 *μ*M, which is 10 to 100 times the maximal amount used in cells (IC50 of 3 to 40 nM)^59^. Stock aliquots of AZD1152 in DMSO were made at 100 mM, so that the final dilution of DMSO in extract was negligible, 1:5000. Fresh egg extract without inhibitor was used as a control. Inhibited and control extracts were imaged simultaneously using multichannel flow chambers made using multiple pieces of double-sided tape.

For inhibiting motors using Na_3_VO_4_ (sodium orthovanadate or ‘vanadate’) (NEB), vanadate was added to extract at a final concentration of 0.5 mM.

*Xenopus laevis* husbandry was done in accordance with the recommendations in the Guide for the Care and Use of Laboratory Animals of the National Institutes of Health. All animals were cared for according to the approved Institutional Animal Care and Use Committee (IACUC) protocol 1941-16 of Princeton University.

### 4-color time lapse TIRFM and image analysis

Total internal reflection fluorescence microscopy (TIRFM) was performed using a Nikon TiE microscope with a 1.49 NA, 100X magnification objective. An Andor Zylsa sCMOS camera was used for acquisition, with software NIS elements (Nikon). Chromatin and kinetochore images were acquired using TIRFM or epifluorescence. For all imaging experiments, multiple fields were imaged in parallel to facilitate sampling more chromosomes. For depletion and drug inhibition experiments, multiple fields were imaged in parallel to enable sampling from experimental and control extract reactions at nearly the same times, and one channel free of chromosomes was imaged to confirm low background nucleation levels in the extract.

The polarity of spindles around chromosomes and the numbers of chromosomes with microtubule networks were determined by manual counting. Normalized tubulin intensity over time was determined by taking the average pixel intensity in a 40 *μ*m x 40 *μ*m window centered around the chromosome over time and dividing by the minimum average value. For measuring intensity over time for the depletion and drug inhibition experiments, a 10 *μ*m x 10 *μ*m window was used to avoid counting microtubules generated from adjacent chromosomes in the same image field. The time of the initial branch was determined by visual inspection. The number of microtubules were counted, and microtubules were tracked using TrackMate v5.2.0 as implemented in Fiji (ImageJ)^60^. The accuracy of the tracking was checked visually.

Plus-end distributions were computed by averaging microtubule tracks into 4 *μ*m bins along the axis of the polar branched network set by the *de novo* mother microtubule. All distributions were then normalized by their maximum values and then averaged together to generate the final distribution.

Chromatin area was determined by thresholding chromatin images in MATLAB such that the intraclass variance between pixel sets was minimized (Otsu’s method), after median filtering. The resulting binarized images were eroded and then dilated. Identical parameters were used for each step while thresholding chromatin images. The locations and angles of the earliest microtubule nucleation events were found manually using Fiji. The number of kinetochores and centromere void regions were determined by visual inspection of kinetochore and chromatin images and tubulin mean intensity time projections.

## 1 Derivation

### 1.1 Dynamic instability

The first step of our theory is to introduce a model for the dynamic instability of microtubules. Following the now standard approach [1], we define *m* (*t, x, ℓ*) (*m_s_*(*t, x, ℓ*)) as the probability density of observing a growing (shrinking) microtubule of length *ℓ* with minus end at *x* at time *t*. We adopt the physiological picture that microtubules grow (shrink) from their plus end at speed *U* (*U_s_*) and are nucleated and stabilized at their minus end. A growing (shrinking) microtubule can stochastically transition into a shrinking (growing) microtubule at a frequency *f*_c_ (*f*_r_) called the catastrophe (rescue) frequency. This entire process occurs far from equilibrium and is regulated by the rate at which GTP-tubulin is hydrolyzed into GDP-tubulin [2]. Because 〈*ℓ*〉/*U* ≪ 〈*ℓ*〉^2^/*D*_M_ is easily satisfied [3], where 〈*ℓ*〉 is the average microtubule length and *D*_M_ is the thermal diffusivity of a microtubule, we may neglect diffusive transport of microtubules. We do not consider active transport by motors in this theory. Therefore, the master equations for the probability densities in the continuum limit are

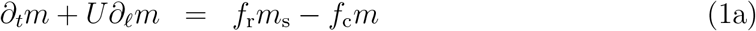

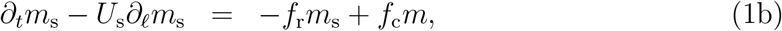

which are of course only valid on a length scale greater than multiple tubulin dimers.

We now make two simplifying assumptions. Firstly, we utilize the fact that rescue events are rare and take *f*_r_ = 0 [4, 5]. That is, once a growing microtubule catastrophes, it will inevitably shrink to zero length. Secondly, we take advantage of the fact that *U_s_* ≫ *U* [6]. That is, microtubules shrink much faster than they grow, and so we take them to effectively disappear once they catastrophe. Under these conditions equations (1) reduce to

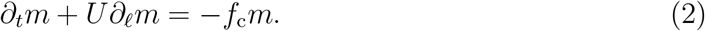

The essential feature of equation (2) is the eventual enforcement of a bounded microtubule length distribution: *m*(*t* → ∞, *x, l*) ~ exp(−*ℓ*/〈*ℓ*〉), where 〈*ℓ*〉 = *U*/*f*_c_ is the average microtubule length. Such exponential distributions have indeed been experimentally verified [3, 7, 8]. Closure of equation (2) requires an initial condition, which we take to be *m*(*t* = 0, *x, ℓ*) = 0, as well as a specification of how new microtubules are nucleated in time and space, which we shall encode in *m*(*t, x, ℓ* = 0).

### 1.2 Microtubule nucleation

We now introduce our model for branching microtubule nucleation regulated by the RanGTP pathway. Branching nucleation, whereby new microtubules nucleate off of pre-existing ones, is catalyzed by proteins effectors that bind to microtubules, some of which (e.g. TPX2 [9–11]) are spatially regulated by the RanGTP pathway and are referred to as a spindle assembly factors (SAFs) [12]. At the onset of spindle assembly, SAFs are sequestered by so-called importin proteins [13]. They are freed to participate in nucleation processes only when RanGTP binds to the importin-SAF complex, which releases the SAF. RanGTP is produced at chromosomes through the RCC1 pathway and released into the cytoplasm, where it can either bind to importin-SAF complexes or be hydrolyzed into its inactive RanGDP form. Thus, RanGTP exists as a gradient around chromosomes.

The simplest model of branching nucleation one can posit is that a branched microtubule is most likely to nucleate wherever the concentration of SAFs bound to pre-existing microtubules is highest. Hence, the probability of nucleating a branched microtubule is proportional to the local concentration of bound SAFs, which we denote by *c*_b_(*t, x*). We note that in reality multiple factors must bind to nucleate a branch [8, 11, 14], some of which are not spatially regulated by RanGTP. However, SAFs bind first [8]. Therefore our model is only strictly valid if the binding of SAFs is rate-limiting. However, taking into account multiple binding events does not affect the main qualitative conclusions of this work [8]. Given these considerations we write the nucleation condition as

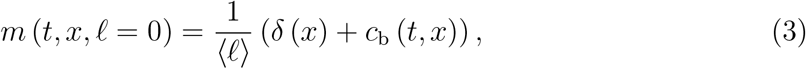

where *δ*(*x*) is the Dirac delta function. The first term captures the initial nucleation of a *de novo* microtubule at *x* = 0 and the second term represents the contribution due to branching nucleation. The prefactor 1/〈*ℓ*〉 sets the units so that 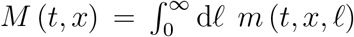 measures the number density of minus ends and 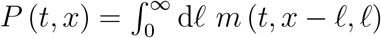 measures the number density of plus ends.

To close the problem, we must express *c*_b_ in terms of m through the reaction-diffusion network regulated by RanGTP (Fig. 3a). It is not our goal here to model the delicate intricacies of the full chemical kinetics involved in the RanGTP pathway [13, 15]. Rather, we seek the simplest tractable description that will preserve the essential microtubule behavior. In this spirit, we first assume the importin binding kinetics are fast enough to come to local equilibrium, which allows us to write the algebraic relation

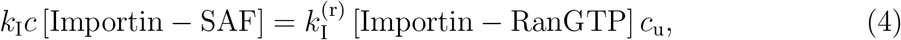

where *c* is the concentration of free RanGTP and *c*_u_ is the concentration of unbound SAFs. Therefore we have that 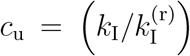 ([Importin − SAF] /[Importin − RanGTP]) *c* ≔ *K*_I_*c*, where we define *K*_I_ to be a constant that encodes all the kinetic activity of the importin molecules. Hence, in this approximation the unbound SAF concentration *c*_u_ is simply proportional to the RanGTP concentration *c*. Together with equation (4), we find that *c* satisfies the reaction-diffusion equation

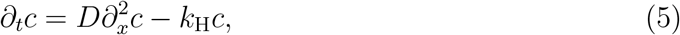

where *D* is the diffusivity of RanGTP and *k*_H_ is the hydrolysis rate of RanGTP → RanGDP. We now assume that protein diffusion is fast compared to microtubule dynamics, i.e. 〈*l*〉^2^/*D* ≪ 〈*l*〉/*U*, so that the RanGTP gradient is quasistatic with respect to the microtubule dynamics [4, 5]. In this limit equation (5) becomes 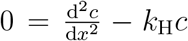. Enforcing the boundary conditions 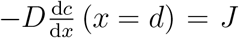 and *c*(|*x*| → ∞) → 0, where *J* is the flux of RanGTP into the cytoplasm produced by chromosomes, furnishes the solution *c*(*x*) = *c*_0_ exp(−|*x* − *d*|/*λ*), where *c*_0_ = *Jλ/D* is the maximum RanGTP concentration at 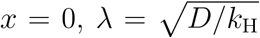 is the characteristic length scale of the RanGTP/SAF gradient, and *d* is the distance between the chromosome and the minus end of the initial *de novo* mother microtubule. Hence the concentration profile of unbound SAFs satisfies

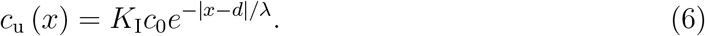

We now obtain the concentration of bound SAFs *c*_b_ needed to close the nucleation condition (3). The simplest way to proceed is to assume that the binding of SAFs to a microtubule occurs at local equilibrium, so that 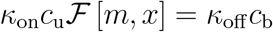, or

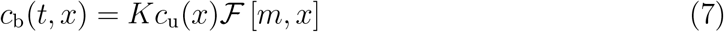

where *K* = *κ*_on_/*κ*_off_ is the ratio between on and off rates of SAFs binding to and unbinding from a microtubule. 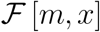 is a functional that expresses the concentration of microtubule lattice at *x* that can serve as a binding site for unbound SAFs at *x*, given the minus end distribution *m*(*t, x′, ℓ*). We can write this functional as

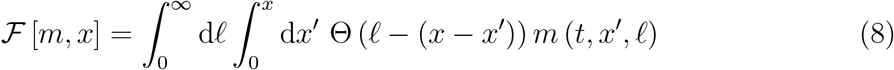

where Θ is the Heaviside step function. The structure of equation (8) ensures that a microtubule with minus end at *x′* can nucleate a branched microtubule at *x* so long as its length *ℓ* is such that *ℓ* > *x* − *x′*. This introduces strong nonlocality to the mathematical formalism and allows us to describe the evolution and variation of the resulting branched network on length scales comparable to and even smaller than the average microtubule length. This generalizes previous continuum theories which were only valid over length scales larger than the average microtubule length [3, 16, 17]. At this stage we have formulated the theory in one spatial dimension, which is justified by the fact that branching angles are shallow, which ensures that the resulting branched networks are highly polar.

### 1.3 Rescaling and problem statement

We now rescale equations (2), (3), and (7) using the dimensionless variables *T* = *tf*_c_, *X* = *x*/〈*l*〉, *L* = *l*/〈*l*〉, and *ψ* = *m*〈*l*〉^2^ to arrive at the dimensionless problem statement

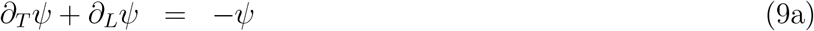

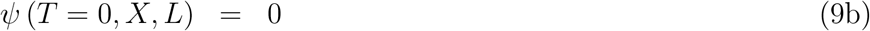

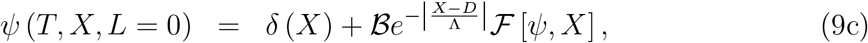

where 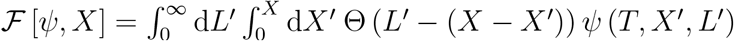. We identify the number 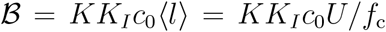 as representing the competition between how many new microtubule branches nucleate versus how many microtubules are lost to catastrophes. Hence we refer to 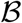 as the *branching number*. Note that in *n* spatial dimensions we would have 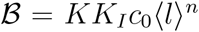. Additionally, two geometric ratios appear. Λ = *λ*/〈*l*〉 is the ratio of the SAF gradient length scale to the average microtubule length, and *D* = *d*/〈*l*〉 is the ratio of the distance between the initial *de novo* nucleation event and the chromosome to the average microtubule length.

Solution of this partial integrodifferential equation for the distribution *ψ* will establish how dynamic microtubules are organized by branching nucleation processes spatially regulated by chromosomes, and hence offer insight into early spindle assembly.

## 2 Branching in a uniform field of SAFs

We first consider the case where the concentration of SAFs is uniform, which corresponds to the limit Λ → ∞ in equations (9). There are two reasons to work in this limit. Firstly, we can perform experiments using a non-hydrolyzable mutant version of RanGTP, RanQ69L, that produces a uniform field of SAFs in *X. laevis* extract, allowing us to benchmark our model on a simpler system (Fig. 2). Secondly, the model has an exact closed-form solution in this limit, which will give us analytical insight into the organization of these dynamic branched networks.

In this limit, the spatial bias imposed by the SAF gradient vanishes. Therefore the kernel of 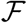 has translational invariance, which motivates solution by Laplace transform. We define 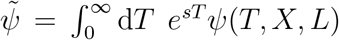. Taking the Laplace transform of equation (9a), using the initial condition (9b), and integrating in *L* gives

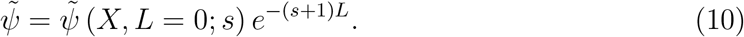

We get 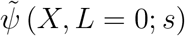 by Laplace transforming the nucleation condition (9c), resulting in the double integral equation

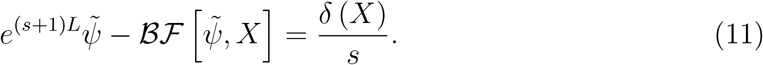

To make progress, we define the spatial Laplace transform of 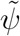 as 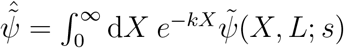. Taking the spatial Laplace transform of equation (11) gives

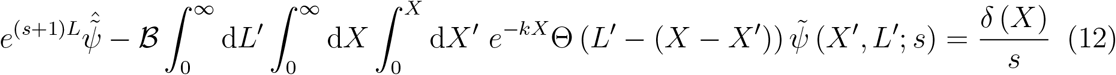

Swapping integration orders on *X* and *X′* factors the spatial integrals into a product of two spatial Laplace transforms in *X′* and *X* − *X′* the latter of which may be explicitly computed. The result is the single integral equation

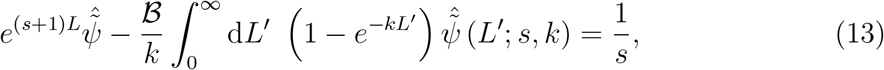

for which the solution may be obtained by proposing the anzats 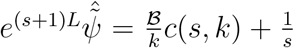 for a to be determined function *c*(*s, k*) [18]. Substituting this anzats into equation (13) and preforming the necessary integrations gives 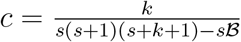, and so we find

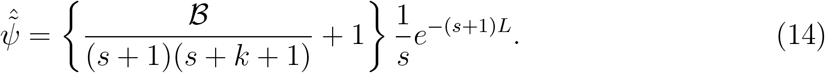

The inverse spatial Laplace transform of equation (14) may be computed and expressed in terms of elementary functions, giving the result

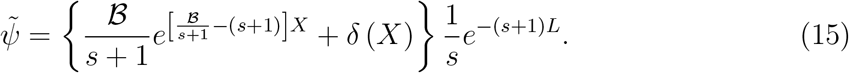

From equation (15) we may directly compute the dimensionless plus end distribution 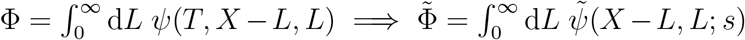, which is the main quantity of interest to us since we can directly track microtubule plus ends. Performing the integration and neglecting the redundant 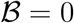 solution gives

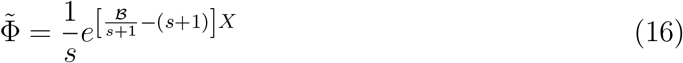

for the plus end distribution. 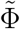 has a simple pole at *s* = 0 and an essential singularity at *s* = −1. Such essential singularities often manifest whenever there are strongly nonlocal physics, such as in the Kosterlitz-Thouless transition [19]. The essential singularity renders the principal part of the local Laurent expansion infinite, precluding the application of standard residue calculus [20]. Therefore, the inverse Laplace transform of equation (16) cannot be computed exactly in terms of known functions, so we employ the Bromwich integral representation

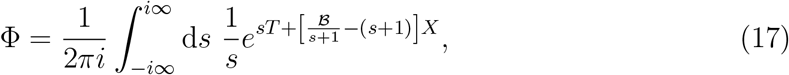

where the notation means we integrate along a vertical line in the complex *s* plane. Note that we have let the integration path tend towards the imaginary axis, since *s* = 0 is just a simple pole, and so the integral is to be interpreted in the principal value sense. Thus, we have derived an exact formula for the plus end distribution of dynamic microtubules branching in a uniform field of SAFs.

We first ask whether the plus end distribution Φ can attain a bounded distribution, which would imply the existence of statistically stationary branched networks. Recall that the final value theorem states 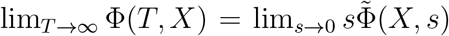 so long as |Φ| is bounded as *T* → ∞. Application of the final value theorem to equation (16) immediately gives

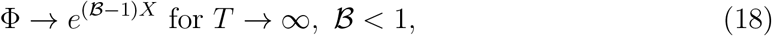

where we require 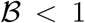 since 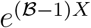 is no longer bounded for all *X* if 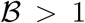. The case where 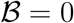 recovers the known result that dynamic microtubules nucleating from a local source have exponentially distributed lengths in the bounded growth regime [1]. The physical interpretation of the distribution (18) is simple: when 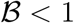, more microtubules are lost to catastrophes than are produced via branching nucleation. Hence, the branched network can only propagate a finite distance before reaching a steady state (Fig. 2d).

The case of 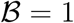 is subtle. The final value theorem (18) would suggest that Φ → 1, but this is not the whole story. Upon numerically evaluating the inverse Laplace transform (17) for 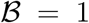 using de Hoog’s method [21, 22], we see that the solution represents traveling waves of constant microtubule density, and therefore no steady state exists (Fig. 2e). In this regime, microtubules are made via branching nucleation as often as they are lost to catastrophes, and so the system can propagate indefinitely while sustaining a constant density. We computed the position of the inflection point of Φ(*T, X*) versus time, *X*\*(*T*), and found a linear relationship, confirming that these profiles indeed correspond to traveling waves. Interestingly, the system selects a wave speed of *V* ≃ *U*/2. Traveling waves of constant microtubule density have been observed in the study of branching asters [16], where the authors provide a similar physical interpretation of the waves, as well as simulations of the traveling microtubule density profiles. Here, we see that their results are a special case of our theory when 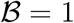, and furthermore, we provide an exact, parameter free formula for such profiles via equation (17).

When 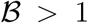, the production of new microtubule branches out competes the loss of microtubules due to catastrophes, and so a branched network forms that propagates outwards and grows autocatalytically with time (Fig. 2f). Interestingly, the position of the maximum of this network, *X*_max_(*T*), also increases linearly with time at a wave speed of *V* ≃ *U*/2. We see that our experimental branched networks fall in this regime for the first ~ 10 min with 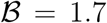 (Fig. 2c). However, after this autocatalytic growth, the experimental branched network saturates in a way consistent with a limiting number of available nucleators.

## 3 Branching in a SAF gradient

We now consider the full problem of microtubules branching in SAF gradient described by equations (9). Because the SAF gradient profile 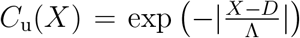 breaks the translational invariance of the problem, an exact closed-form solution is no longer possible. Furthermore, regular perturbation methods are ill-suited for systems that exhibit autocatalytic growth. Hence, we proceed to solve the complete equations (9) numerically.

We begin by working with the Laplace transformed distribution 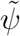. In the case where there is a SAF gradient, equation (11) becomes

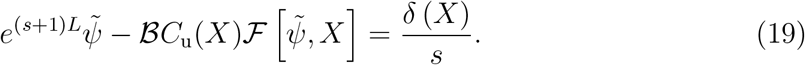

We now substitute the ansatz

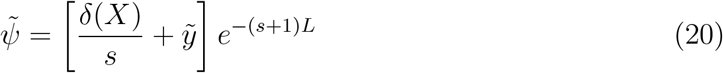

into equation (19) for a to be determined function 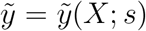. Evaluating the integrals appear upon this substitution results in

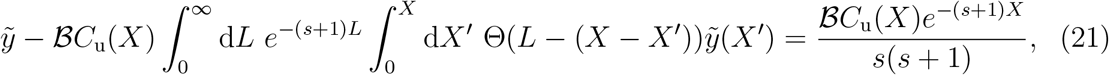

which is an integral equation for 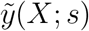. In order to solve equation (21) numerically, we discretize all variables onto a regular grid *X* → *X_i_* and 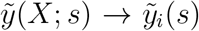 for *i* = 0,…, *M* and we choose Δ*X* = *X*_*i*+1_ − *X_i_* = 0.1. We first evaluate the integral over *X′* using the trapezoidal rule and then evaluate the integral over *L* analytically. The result is

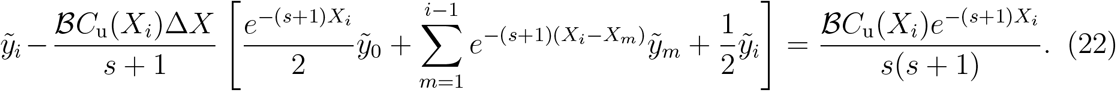

Equation (22) has the structure of the linear system 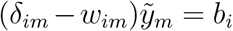 (Einstein summation convention is used here), which we invert using standard LU decomposition to find the desired 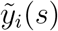.

With 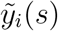 now constructed, we numerically compute the inverse Laplace transform using de Hoog’s method [21, 22] to generate *y_i_*(*T*), which allows us to compute *ψ*(*T, X_i_, L*) using equation (20). The desired plus end distribution is computed from 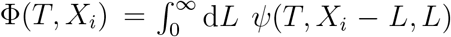 using linear interpolation and the trapezoidal rule. We benchmarked our numerical method using *C*_u_(*X*) = 1 to ensure that it reproduced the results of the uniform field branched networks.

We find that the parameter choices of 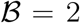 and Λ = 3 best reproduce the experimental branched networks that form around chromosomes (Fig. 3b).

## REFERENCES

1. McKim, K. S. & Hawley, R. S. Chromosomal Control of Meiotic Cell Division. Science (1995) doi:10.1126/science.270.5242.1595.

2. Ohkura, H. Meiosis: An Overview of Key Differences from Mitosis. Cold Spring Harb Perspect Biol 7, a015859 (2015).

3. Dumont, J. & Desai, A. Acentrosomal spindle assembly and chromosome segregation during oocyte meiosis. Trends in Cell Biology 22, 241–249 (2012).

4. Zhang, H. & Dawe, R. K. Mechanisms of plant spindle formation. Chromosome Res 19, 335–344 (2011).

5. Karsenti, E., Newport, J. & Kirschner, M. Respective roles of centrosomes and chromatin in the conversion of microtubule arrays from interphase to metaphase. Journal of Cell Biology 99, 47s–54s (1984).

6. Heald, R. et al. Self-organization of microtubules into bipolar spindles around artificial chromosomes in Xenopus egg extracts. Nature 382, 420–425 (1996).

7. Dinarina, A. et al. Chromatin Shapes the Mitotic Spindle. Cell 138, 502–513 (2009).

8. McGill, M. & Brinkley, B. R. Human chromosomes and centrioles as nucleating sites for the in vitro assembly of microtubules from bovine brain tubulin. Journal of Cell Biology 67, 189–199 (1975).

9. Witt, P. L., Ris, H. & Borisy, G. G. Origin of kinetochore microtubules in Chinese hamster ovary cells. Chromosoma 81, 483–505 (1980).

10. De Brabander, M., Geuens, G., Mey, J. D. & Joniau, M. Nucleated assembly of mitotic microtubules in living PTK2 cells after release from nocodazole treatment. Cell Motility 1, 469–483 (1981).

11. Khodjakov, A., Copenagle, L., Gordon, M. B., Compton, D. A. & Kapoor, T. M. Minus-end capture of preformed kinetochore fibers contributes to spindle morphogenesis. Journal of Cell Biology 160, 671–683 (2003).

12. Maiato, H., Rieder, C. L. & Khodjakov, A. Kinetochore-driven formation of kinetochore fibers contributes to spindle assembly during animal mitosis. Journal of Cell Biology 167, 831–840 (2004).

13. Kitamura, E. et al. Kinetochores Generate Microtubules with Distal Plus Ends: Their Roles and Limited Lifetime in Mitosis. Developmental Cell 18, 248–259 (2010).

14. Mishra, R. K., Chakraborty, P., Arnaoutov, A., Fontoura, B. M. A. & Dasso, M. The Nup107-160 complex and γ-TuRC regulate microtubule polymerization at kinetochores. Nat Cell Biol 12, 164–169 (2010).

15. Sikirzhytski, V. et al. Microtubules assemble near most kinetochores during early prometaphase in human cells. Journal of Cell Biology 217, 2647–2659 (2018).

16. Cavazza, T. & Vernos, I. The RanGTP Pathway: From Nucleo-Cytoplasmic Transport to Spindle Assembly and Beyond. Frontiers in Cell and Developmental Biology 3, (2016).

17. Carazo-Salas, R. E. et al. Generation of GTP-bound Ran by RCC1 is required for chromatin-induced mitotic spindle formation. Nature 400, 178–181 (1999).

18. Kalab, P., Pu, R. T. & Dasso, M. The Ran GTPase regulates mitotic spindle assembly. Current Biology 9, 481–484 (1999).

19. Ohba, T., Nakamura, M., Nishitani, H. & Nishimoto, T. Self-Organization of Microtubule Asters Induced in Xenopus Egg Extracts by GTP-Bound Ran. Science (1999) doi:10.1126/science.284.5418.1356.

20. Wilde, A. & Zheng, Y. Stimulation of Microtubule Aster Formation and Spindle Assembly by the Small GTPase Ran. Science (1999) doi:10.1126/science.284.5418.1359.

21. Zhang, C., Hughes, M. & Clarke, P. R. Ran-GTP stabilises microtubule asters and inhibits nuclear assembly in Xenopus egg extracts. Journal of Cell Science 112, 2453–2461 (1999).

22. Kalab, P., Weis, K. & Heald, R. Visualization of a Ran-GTP Gradient in Interphase and Mitotic Xenopus Egg Extracts. Science (2002) doi:10.1126/science.1068798.

23. Kaláb, P., Pralle, A., Isacoff, E. Y., Heald, R. & Weis, K. Analysis of a RanGTP-regulated gradient in mitotic somatic cells. Nature 440, 697–701 (2006).

24. Gruss, O. J. et al. Ran Induces Spindle Assembly by Reversing the Inhibitory Effect of Importin α on TPX2 Activity. Cell 104, 83–93 (2001).

25. Petry, S., Groen, A. C., Ishihara, K., Mitchison, T. J. & Vale, R. D. Branching Microtubule Nucleation in Xenopus Egg Extracts Mediated by Augmin and TPX2. Cell 152, 768–777 (2013).

26. David, A. F. et al. Augmin accumulation on long-lived microtubules drives amplification and kinetochore-directed growth. Journal of Cell Biology 218, 2150–2168 (2019).

27. Decker, F., Oriola, D., Dalton, B. & Brugués, J. Autocatalytic microtubule nucleation determines the size and mass of Xenopus laevis egg extract spindles. eLife 7, e31149 (2018).

28. Carmena, M., Wheelock, M., Funabiki, H. & Earnshaw, W. C. The chromosomal passenger complex (CPC): from easy rider to the godfather of mitosis. Nat Rev Mol Cell Biol 13, 789–803 (2012).

29. Sampath, S. C. et al. The Chromosomal Passenger Complex Is Required for Chromatin-Induced Microtubule Stabilization and Spindle Assembly. Cell 118, 187–202 (2004).

30. Maresca, T. J. et al. Spindle Assembly in the Absence of a RanGTP Gradient Requires Localized CPC Activity. Current Biology 19, 1210–1215 (2009).

31. Loughlin, R., Heald, R. & Nédélec, F. A computational model predicts Xenopus meiotic spindle organization. Journal of Cell Biology 191, 1239–1249 (2010).

32. Brugués, J., Nuzzo, V., Mazur, E. & Needleman, D. J. Nucleation and Transport Organize Microtubules in Metaphase Spindles. Cell 149, 554–564 (2012).

33. Oh, D., Yu, C.-H. & Needleman, D. J. Spatial organization of the Ran pathway by microtubules in mitosis. PNAS 113, 8729–8734 (2016).

34. Wollman, R. et al. Efficient Chromosome Capture Requires a Bias in the ‘Search-and-Capture’ Process during Mitotic-Spindle Assembly. Current Biology 15, 828–832 (2005).

35. Paul, R. et al. Computer simulations predict that chromosome movements and rotations accelerate mitotic spindle assembly without compromising accuracy. PNAS 106, 15708–15713 (2009).

36. Petry, S., Pugieux, C., Nédélec, F. J. & Vale, R. D. Augmin promotes meiotic spindle formation and bipolarity in Xenopus egg extracts. PNAS 108, 14473–14478 (2011).

37. Lüders, J., Patel, U. K. & Stearns, T. GCP-WD is a γ-tubulin targeting factor required for centrosomal and chromatin-mediated microtubule nucleation. Nat Cell Biol 8, 137–147 (2006).

38. Thawani, A., Stone, H. A., Shaevitz, J. W. & Petry, S. Spatiotemporal organization of branched microtubule networks. eLife 8, e43890 (2019).

39. Ishihara, K., Korolev, K. S. & Mitchison, T. J. Physical basis of large microtubule aster growth. eLife 5, e19145 (2016).

40. Reber, S. B. et al. XMAP215 activity sets spindle length by controlling the total mass of spindle microtubules. Nat Cell Biol 15, 1116–1122 (2013).

41. Gaglio, T. et al. Opposing motor activities are required for the organization of the mammalian mitotic spindle pole. Journal of Cell Biology 135, 399–414 (1996).

42. Schuh, M. & Ellenberg, J. Self-Organization of MTOCs Replaces Centrosome Function during Acentrosomal Spindle Assembly in Live Mouse Oocytes. Cell 130, 484–498 (2007).

43. Sluder, G. & Rieder, C. L. Centriole number and the reproductive capacity of spindle poles. Journal of Cell Biology 100, 887–896 (1985).

44. Wang, S. Z. & Adler, R. Chromokinesin: a DNA-binding, kinesin-like nuclear protein. Journal of Cell Biology 128, 761–768 (1995).

45. Molina, I. et al. A Chromatin-associated Kinesin-related Protein Required for Normal Mitotic Chromosome Segregation in Drosophila. Journal of Cell Biology 139, 1361–1371 (1997).

46. Ruden, D. M., Cui, W., Sollars, V. & Alterman, M. ADrosophilaKinesin-like Protein, Klp38B, Functions during Meiosis, Mitosis, and Segmentation. Developmental Biology 191, 284–296 (1997).

47. Alphey, L. et al. KLP38B: A Mitotic Kinesin-related Protein That Binds PP1. Journal of Cell Biology 138, 395–409 (1997).

48. Schneider, M. W. G. et al. A chromatin phase transition protects mitotic chromosomes against microtubule perforation. 2021.07.05.450834 https://www.biorxiv.org/content/10.1101/2021.07.05.450834v1 (2021) doi:10.1101/2021.07.05.450834.

49. Dalton, B. A., Oriola, D., Decker, F., Jülicher, F. & Brugués, J. A gelation transition enables the self-organization of bipolar metaphase spindles. 2021.01.15.426844 https://www.biorxiv.org/content/10.1101/2021.01.15.426844v1 (2021) doi:10.1101/2021.01.15.426844.

50. King, M. & Petry, S. Visualizing and Analyzing Branching Microtubule Nucleation Using Meiotic Xenopus Egg Extracts and TIRF Microscopy. in The Mitotic Spindle (eds. Chang, P. & Ohi, R.) vol. 1413 77–85 (Springer New York, 2016).

51. Gell, C. et al. Chapter 13 - Microtubule Dynamics Reconstituted In Vitro and Imaged by Single-Molecule Fluorescence Microscopy. in Methods in Cell Biology (eds. Wilson, L. & Correia, J. J.) vol. 95 221–245 (Academic Press, 2010).

52. Gasser, S. M. & Laemmli, U. K. Improved methods for the isolation of individual and clustered mitotic chromosomes. Experimental Cell Research 173, 85–98 (1987).

53. Sone, T. et al. Changes in Chromosomal Surface Structure by Different Isolation Conditions. Archives of Histology and Cytology 65, 445–455 (2002).

54. Maeshima, K. & Laemmli, U. K. A Two-Step Scaffolding Model for Mitotic Chromosome Assembly. Developmental Cell 4, 467–480 (2003).

55. Fukui, K., Takata, H. & Uchiyama, S. Preparation Methods of Human Metaphase Chromosomes for their Proteome Analysis. in Organelle Proteomics (eds. Pflieger, D. & Rossier, J.) 149–160 (Humana Press, 2008). doi:10.1007/978-1-59745-028-7_10.

56. Ma, H. T. & Poon, R. Y. C. Synchronization of HeLa Cells. in Cell Cycle Synchronization: Methods and Protocols (ed. Banfalvi, G.) 151–161 (Humana Press, 2011). doi:10.1007/978-1-61779-182-6_10.

57. Murray, A. W. & Kirschner, M. W. Cyclin synthesis drives the early embryonic cell cycle. Nature 339, 275–280 (1989).

58. Hannak, E. & Heald, R. Investigating mitotic spindle assembly and function in vitro using Xenopus laevis egg extracts. Nat Protoc 1, 2305–2314 (2006).

59. Yang, J. et al. AZD1152, a novel and selective aurora B kinase inhibitor, induces growth arrest, apoptosis, and sensitization for tubulin depolymerizing agent or topoisomerase II inhibitor in human acute leukemia cells in vitro and in vivo. Blood 110, 2034–2040 (2007).

60. Tinevez, J.-Y. et al. TrackMate: An open and extensible platform for single-particle tracking. Methods 115, 80–90 (2017).

## References

[1] Marileen Dogterom and Stanislas Leibler. Physical aspects of the growth and regulation of microtubule structures. Physical review letters, 70(9):1347, 1993.

[2] Thomas CT Michaels, Shuo Feng, Haiyi Liang, and L Mahadevan. Mechanics and kinetics of dynamic instability. Elife, 9:e54077, 2020.

[3] Franziska Decker, David Oriola, Benjamin Dalton, and Jan Brugués. Autocatalytic microtubule nucleation determines the size and mass of xenopus laevis egg extract spindles. Elife, 7:e31149, 2018.

[4] R Wollman, EN Cytrynbaum, JT Jones, T Meyer, JM Scholey, and A Mogilner. Efficient chromosome capture requires a bias in the ‘search-and-capture’ process during mitotic-spindle assembly. Current Biology, 15(9):828–832, 2005.

[5] Raja Paul, Roy Wollman, William T Silkworth, Isaac K Nardi, Daniela Cimini, and Alex Mogilner. Computer simulations predict that chromosome movements and rotations accelerate mitotic spindle assembly without compromising accuracy. Proceedings of the National Academy of Sciences, 106(37):15708–15713, 2009.

[6] Christopher Gell, Volker Bormuth, Gary J Brouhard, Daniel N Cohen, Stefan Diez, Claire T Friel, Jonne Helenius, Bert Nitzsche, Heike Petzold, Jan Ribbe, et al. Microtubule dynamics reconstituted in vitro and imaged by single-molecule fluorescence microscopy. In Methods in cell biology, volume 95, pages 221–245. Elsevier, 2010.

[7] Jan Brugués, Valeria Nuzzo, Eric Mazur, and Daniel J Needleman. Nucleation and transport organize microtubules in metaphase spindles. Cell, 149(3):554–564, 2012.

[8] Akanksha Thawani, Howard A Stone, Joshua W Shaevitz, and Sabine Petry. Spatiotemporal organization of branched microtubule networks. eLife, 8:e43890, 2019.

[9] Raymundo Alfaro-Aco, Akanksha Thawani, and Sabine Petry. Biochemical reconstitution of branching microtubule nucleation. eLife, 9, 2020.

[10] Matthew R King and Sabine Petry. Phase separation of tpx2 enhances and spatially coordinates microtubule nucleation. Nature Communications, 11(1):1–13, 2020.

[11] Sagar U Setru, Bernardo Gouveia, Raymundo Alfaro-Aco, Joshua W Shaevitz, Howard A Stone, and Sabine Petry. A hydrodynamic instability drives protein droplet formation on microtubules to nucleate branches. Nature Physics, 17(4):493–498, 2021.

[12] Petr Kalab and Rebecca Heald. The rangtp gradient-a gps for the mitotic spindle. Journal of cell science, 121(10):1577–1586, 2008.

[13] Maïwen Caudron, Gertrude Bunt, Philippe Bastiaens, and Eric Karsenti. Spatial coordination of spindle assembly by chromosome-mediated signaling gradients. Science, 309(5739):1373–1376, 2005.

[14] Sabine Petry, Aaron C Groen, Keisuke Ishihara, Timothy J Mitchison, and Ronald D Vale. Branching microtubule nucleation in xenopus egg extracts mediated by augmin and tpx2. Cell, 152(4):768–777, 2013.

[15] Chaitanya A Athale, Ana Dinarina, Maria Mora-Coral, Celine Pugieux, Francois Nedelec, and Eric Karsenti. Regulation of microtubule dynamics by reaction cascades around chromosomes. Science, 322(5905):1243–1247, 2008.

[16] Keisuke Ishihara, Kirill S Korolev, and Timothy J Mitchison. Physical basis of large microtubule aster growth. Elife, 5:e19145, 2016.

[17] Bryan Kaye, Olivia Stiehl, Peter J Foster, Michael J Shelley, Daniel J Needleman, and Sebastian Fürthauer. Measuring and modeling polymer concentration profiles near spindle boundaries argues that spindle microtubules regulate their own nucleation. New Journal of Physics, 20(5):055012, 2018.

[18] Subrahmanyan Chandrasekhar. Radiative transfer. Courier Corporation, 2013.

[19] John Michael Kosterlitz and David James Thouless. Ordering, metastability and phase transitions in two-dimensional systems. Journal of Physics C: Solid State Physics, 6(7):1181, 1973.

[20] George B Arfken and Hans J Weber. Mathematical methods for physicists. American Association of Physics Teachers, 1999.

[21] Frank R De Hoog, JH Knight, and AN Stokes. An improved method for numerical inversion of laplace transforms. SIAM Journal on Scientific and Statistical Computing, 3(3):357–366, 1982.

[22] Fredrik Johansson et al. mpmath: a Python library for arbitrary-precision floating-point arithmetic (version 0.18), December 2013. http://mpmath.org/.

